# The complex molecular basis of enhanced stress resilience in extreme drought-tolerant *Arabis* grassland species

**DOI:** 10.1101/2025.10.02.680103

**Authors:** Abdul Saboor Khan, Simon Maria Zumkeller, Gregor Schmitz, Irina Calic, Tahir Ali, Neda Rahnamae, Lea Hoerdemann, Lina Abdelwahed, Jedrzej Szymanski, Juliette de Meaux

## Abstract

**Background and Aims:** Plant species in competitive meadows must tolerate extreme stress, yet the mechanisms underlying resilience remain poorly understood. *Arabis nemorensis*, an endangered selfing species of Euro- pean floodplain grasslands, experiences both flooding and drought and hybridizes with its close relative, *A. sagittata*. We investigated how these species differ in drought survival and the molec- ular basis of their responses.

**Methods:** Sympatric lineages of *A. nemorensis* and *A. sagittata* were compared in a controlled dry-down experiment, complemented by transcriptome and small RNA profiling, and machine-learning anal- ysis of cis-regulatory motifs.

**Key Results:** Both species wilted at 5% soil moisture, but *A. sagittata* recovered more effectively (90% vs. 50%). This difference was not explained by a major QTL, suggesting a polygenic basis. Transcrip- tome profiling revealed stronger induction in *A. sagittata* (6,359 vs. 5,571 differentially expressed genes). Small RNA analysis identified species-specific regulation of miR408, a conserved drought regulator. Machine-learning identified 307 sequence motifs predictive of stress-responsive expres- sion, with motif distributions indicating distinct regulatory networks.

**Conclusions:** This study reveals the polygenic and regulatory complexity underlying divergent drought resili- ence strategies in the closely related species thriving in grassland environments.

*Arabis nemorensis* and its close relative *A. sagittata* co-occur in a floodplain meadow exposed to flooding and drought. In dry-down experiments, *A. sagittata* recovered more effectively than *A. nemorensis*. Transcriptome and small RNA analyses revealed stronger stress responses in *A. sagittata*, including regulation of miR408. These differences result from different regulatory networks and have a polygenic basis.

## Introduction

In response to rapid climate change, plants are expected to be challenged by episodes of increasingly severe and frequent drought, threatening global food security and natural ecosystems (Carrão et al., 2016). The need to create high-yield cultivars that use water more efficiently than their current counterparts is pressing (Onyekachi et al., 2019; Gupta et al., 2020). Physiological and molecular changes due to drought stress have been widely studied in crops and model plant species, yet which of these changes determines survival to natural drought events is poorly understood (Bresson et al., 2013; Rizhsky et al., 2004; Pellegrineschi et al., 2004). Global patterns of plant forms and strategies indicate that tolerance to challenging environmental conditions has evolved as a trade-off with growth rates (Grime, 1977; Díaz et al., 2007). Plant species that have evolved to balance the intense competition that characterizes dense communities with the demands imposed by abiotic stress such as plants that have adapted to dry grasslands may reveal how these plants evolve the ability to tolerate.

Drought stress experienced in dry grasslands pose particularly acute challenges to plant growth, because adapted species must not only survive but also prevail in the dense and competitive communities that host them (Joyce et al., 2016; da Silva et al., 2013; Kübert et al., 2019). Dry grassland species have evolved sophisticated adaptations to cope with water scarcity, such as deep root systems, tissues that store water, or leaf structures that tolerate drought (Májeková et al., 2019). However, prolonged drought can exceed the limits of these adaptations, leading to reduced plant growth, productivity, or survival (Joyce et al., 2016; Lei et al., 2016). Drought stress can also exacerbate the effects of other environmental stresses, such as nutrient loading and pollution (Kübert et al., 2019). On the other hand, in grasslands exposed to intense drought stress, drought tolerance may improve resistance to competition (Mount et al., 2023). The better we understand the physiological and molecular responses of grassland species to drought stress, the more we can hope to enhance the resilience of grassland ecosystems to environmental challenges (da Silva et al., 2013; Lei et al., 2016, Rhee et al., 2024).

Drought stress activates many physiological responses (Bohnert et al., 1995; Wang et al., 2021; Xiong and Zhu, 2002). Although the early events in the perception of stress signals are unclear, the effects of drought stress on gene expression are not: in plants they are global and massive (Zhang et al., 2018). Some physiological changes are regulated at the transcriptional level (Wilkins et al., 2010; Baerenfaller et al., 2012; Hura et al., 2022; Harb et al., 2010; Shinozaki and Yamaguchi-Shinozaki, 2007; Zhang et al., 2018). Many genes involved in osmotic adjustment, stress signaling, and antioxidant defense are up-regulated while genes involved in growth and development are downregulated (Yang et al., 2021). Alterations of protein activity and metabolite abundance also contribute to mechanisms that defend against drought (Shinozaki and Yamaguchi-Shinozaki, 2007; Ozfidan et al., 2012; Cruz de Carvalho, 2008): amendments to pigment composition and photosynthesis, for example, are among the most critical responses required (Liu et al., 2022).

Few studies investigate the genetic basis of drought tolerance in species adapted to extreme environment. Unfortunately, the slow growth of most drought-tolerant species limits our ability to dissect the genetic basis of their capacity to tolerate low water levels (Anithakumari et al., 2012; Lopes et al., 2011). However, the *Arabis* genus has several biannual species that grow in competitive grassland meadows. *Arabis nemonresis,* for example, grows exclusively in floodplain grasslands, where it can withstand both protracted submersion during flooding episodes and severe summer droughts (Hölzel and Otte, 2001). A close relative, *Arabis sagittata,* grows in dry calcareous grasslands across South to Central Europe (Karl and Koch, 2014; Titz, 1972). The low genetic variability of these species (Dittberner et al., 2019) makes their extinction likely, especially as natural grassland habitats are rapidly shrinking. Interestingly, the two species have exchanged alleles via gene flow in the past, and gene flow has been documented today in certain populations (Dittberner et al., 2022).

Do the grassland species *A. nemorensis* and *A. sagittata* differ in their tolerance to drought? Our data show that both species tolerate extremely low levels of soil water content (SWC), but, as their ecological preferences suggest, *A. sagittata* survives wilting better than the floodplain- specific species *A. nemorensis*. We investigated the variation in gene expression associated with these differences. Compared to *A. nemorensis,* the more drought-tolerant *A. sagittata* shows an increase of stress-related genes and a preferential reactivation of growth-related genes. Furthermore, responses of both species could be related to different sets of DNA sequence motifs, suggesting transcription factors play a role. Ultimately, given the differential activation of microRNA408 in *A. sagittata,* we conclude that post-transcriptional regulation also contribute to drought tolerance in this system.

## Materials and Methods

### Plant materials and growth conditions

For the dry-down experiments, seeds of *A. nemorensis,* genotype 10, and *A. sagittata,* genotype 69, were grown in 100 completely randomized replicates. These two genotypes were collected in Riedstadt (Hessen, Germany) by Dittberner et al. (2019). Both genotypes originate from a floodplain grassland site where *A. nemorensis,* and *A. sagittata* occur in sympatry. *A. nemorensis* is found primarily on floodplains, whereas *A. sagittata* is found mostly in dry calcareous grasslands (Karl & Koch, 2014; Hand & Gregor, 2006). For the QTL mapping experiments, we grew one offspring of each of 300 F2 individuals, obtained by crossing genotypes 10 and 69 (Rahnamae et al., 2025). Each individual plant was genotyped following protocols described in Rahnamae et al. (2025).

### Dry-down experimental protocol and phenotypic measurements

The dry-down experiment was conducted following the methodology described in Bouzid et al. (2019). After a period of acclimation at 60% SWC, watering was stopped, and SWC was quantified every day until wilting using the precision balance. The day plants showed the first symptoms of wilting was scored. The plants were re-watered 4 days after wilting, and kept watering two times per week once they started producing fresh leaves.

### Phenotypic trait measurement

Phenotypic differences between the two species were assessed from the first day of water withdrawal until wilting and during recovery. In total, 10 phenotypes were measured throughout the dry-down experiment: initial rosette area, initial leaf thickness and leaf thickness at wilting, daily SWC, rate of water loss, rate of recovery, time to recovery, and severity of damage. In addition, we measured leaf area of three medium-sized leaves for 10 plants of each species as well as stomata length and stomata density (**details in Suppl. Methods**).

### Statistical analysis of the phenotypic data

Data visualization was performed using the CRAN package ggplot2 (Wickham, 2009). Statistical differences between the two species were assessed using a generalized linear model (GLM) in R. All models included block as a factor. For traits monitored over time, time was incorporated to assess differences in rates. Error distributions were specified based on the nature of the phenotypic trait. A binomial family of GLMs was applied to traits such as recovery, whereas a quasipoisson distribution was used for all other phenotypic traits to account for overdispersion.

### Analysis of transcriptome and small RNA variation during dry-down

To quantify transcript abundance during drought stress, leaf samples were collected for RNA extraction. Depending on leaf size, one or half of a young leaf was sampled from four biological replicates per species at three time points: (1) control plants (60% SWC), (2) wilting plants (5% SWC), and (3) leaves formed during the recovery phase (10-15 days post re-watering). All samples were flash-frozen in liquid nitrogen and stored at -80°C in tubes containing 10 metal beads, and RNA was extracted **(details in Suppl. Methods)**. For small RNA sequencing, leaf samples were collected following the protocol described above. Small RNA was extracted using the Qiagen RNeasy Plant Mini Kit (Qiagen, Germany). A total of 24 leaf RNA samples were sequenced using Illumina SE50 technology at the Cologne Center for Genomics (CCG), Germany (**details in Suppl. Methods**).

### Bioinformatics analysis of RNA transcriptome

Transcriptome data were first trimmed and filtered using the FastQC package (v0.11.4). We then used Hisat2 to map the trimmed and filtered reads to the *A. nemonresis* high-quality reference genome with reference PRJEB8986 in ENA (Rahnamae et al., 2025). Using Samtools (version 1.3.1), only high-quality, uniquely mapped reads with a correct mapping probability of ≥ 90% were kept, resulting in on average 86% of the reads. RNA integrity was verified using a custom R script to confirm uniform transcript coverage and ensure that RNA degradation did not bias expression estimates. Gene expression was quantified using DESeq2 (bioconductor version: release 3.5) to identify differentially expressed genes (DEGs) (Love et al., 2014). Contrasts were applied to identify DEGs in both *A. nemorensis* and *A. sagittata* across all three conditions **(details on package parameters and scripts are given in Suppl. Methods)**.

### Identification of expression predictive motifs (EPMs)

To explore whether we could identify regulatory motifs associated with differential gene expression, we used DeepCRE (Peleke et al., 2024; Zumkeller et al., 2025). This approach uses a machine-learning strategy to identify the features of cis-regulatory sequences that predict whether a gene is going to be strongly or weakly expressed under any conditions and in all species. Cis-regulatory sequences were considered to be 1000 nt upstream of the gene’s transcription start site (TSS), 1000 nt downstream of the gene’s transcription termination site (TTS), 500 nt downstream from the transcription start site (TSS), and 500 nt upstream from the transcription termination site (TTS). Genes with high transcript profiles were labeled as such if these belonged to the upper 20th percentile of the log-median of transcript per million. The same number of genes was sampled from a pool of genes with a log-median of zero or minimally larger than zero. Accordingly, the classes were balanced, and percentile threshold were found to be similar across different species and under various conditions. Models were trained on each chromosome separately (**details on model training are given in Suppl. Methods**).

Using the genes correctly predicted by these models, we extracted importance scores of each nucleotide position and each sequence motif using SHAPley and TF-MoDisco (Lundberg and Lee, 2017; Shrikumar et al., 2018). In the context of this analysis, Shapley values quantify the importance of each nucleotide position or sequence motif in predicting the expression class of the gene. Saliency maps then describe variation in nucleotide importance as a function of the distance to the TSS and to the TTS, respectively. Based on similarity in both sequence and importance, expression predicting motifs (EPMs) were assembled using TF-MoDiSCO (Shrikumar et al., 2018). EPMs, like transcription factor binding sites, are described by contribution weight matrices (CWMs), but the CWM of transcription factor binding sites describes whether a sequence occurs or not in the DNA, the CWM of EPMs capture the importance of nucleotides for gene expression prediction. EPMs are thus assigned separately for each species, each condition, and positional preference relative to the TSS and TTS. In addition, some EPMs predict high rates of expression, others low rates.

The occurrence of EPMs was compared across species and treatments with a chi-squared test of independence. To identify shared EPMs across species and treatments, we measured EPM similarity using the Sandelin-Wasserman algorithm and divided the results into 25 clusters using UMAP in R (Wasserman and Sandelin, 2004; Dalmia and Sia, 2021). For determination of the number of clusters we followed Peleke et al. (2024), except that we used the next minimum from Elbow and Silhouette methods of cluster EPMs to identify the best clustering for *A. nemorensis* and *A. sagittata.* We then analyzed which EPM clusters were predictive of expression under different treatments within the same species or between the two species for each treatment.

### GO term enrichment analysis

*A. thaliana* orthologs of the annotated genes in *A. nemorensis* or *A. sagittata* were identified via Orthofinder, and genes lacking orthologs were excluded from this analysis (Emms and Kelly, 2019). The comprehensive set of unique orthologs expressed in our dataset served as the background universe for gene ontology (GO) enrichment analysis. GO enrichment analysis was conducted using the topGO (2.59.0) package in R (Alexa, Rahnenfuhrer and Lengauer, 2006). Emphasis was placed on the GO terms related to biological processes, employing the ‘elim’ algorithm alongside Fisher’s exact test to account for GO hierarchy, using org.At.tair.db (3.21.0) for annotations. GO terms with a Fisher *p* below 0.0001 were deemed statistically significant (following He et al., 2016). Since genes sharing EPMs are predicted to be co-regulated, we also used GO enrichment analyses to gain further insight into the molecular functions of genes sharing EPMs. We focused on the EPMs extracted from the response to wilting and mapped their occurrence in the genome using BLAMM (Fostier, 2020), then filtered these using criteria analogous to those in Peleke et al. (2024) (BLAMM score >10) onto the annotated genomes of *A. nemorensis* and *A. sagittata*.

### Analysis of regulatory elements in *A. nemorensis* and *A. sagittata* miR408 alleles

Flanking sequences (1 kbp up- and downstream) of *A. thaliana* miRNA408 (At; LR797788.1: 19847354 - 19849574) were retrieved from TAIR10.v55. These sequences were then used to identify homologous regions in the genomes of *A. nemorensis* (An; chr4: 41024454 - 41029450) and *A. sagittata* (As; chr4: 42812501 - 42817531) using BLASTn (Z. Zhang et al., 2000). The identified *A. nemorensis* and *A. sagittata* sequences were subsequently aligned with AtmiRNA408 using MAFFT and visualized with AliView (Katoh, 2002; Larsson, 2014), resulting in a 2038 bp alignment. To understand the regulatory mechanisms of miRNA408-5p, we mapped EPMs from all treatments onto this alignment. This was combined with predicting potential transcription factor binding site (TFBS) using PCBase 2.0 data from *A. thaliana* ChIP-seq experiments available in the PlantPAN tool (Chow et al., 2019) .

Additionally, we investigated if potential targets of miRNA408-5p were recognized by DeepCRE models. Using blastn, we also identified a potential target gene with differential regulation under wilting conditions in *A. nemorensis* (An; chr6:12870083 - 12875969) and *A. sagittata* (As; chr6:17611380 - 17617190) and determined its EPM.

## Results

### Higher survival rates in response to wilting in *A. sagittata* compared to in *A. nemorensis*

We recorded the onset of wilting symptoms after watering was stopped and found that *A. nemorensis* and *A. sagittata* plants did not differ significantly in the number of days until wilting (F_1,197_ = 2.5736, *p* = 0.239, **Figure 1A-D, Table-S1, Fig-S1**). Both species wilted at a remarkably low SWC of <5% (*A. sagittata* 4.16% and *A. nemorensis* 3.53%) (**Fig-S2A**). Rosette size was significantly larger in *A. nemorensis* size than in *A. sagittata* (47.837 cm^2^ vs. 37.439 cm^2^, F_1,197_ = 32.5540, *p* = 9.26e-08, **Figure 1E)**, suggesting that larger leaf surface area resulted in more water loss. Independent of species, the decrease in SWC was significantly correlated with days to wilting (F_1,197_ = 21.608, *p* = 1.25e-12, **Fig-S2B**, and F_1,1364.2 =_ 0.4998, *p* = 0.4799, **Fig-S3**). The interaction between species and initial rosette area had a significant effect on the number of days until wilting (F_1,195_ = 1.1991, *p* = 0.00804, **Figure 1F**). Desiccation rate, measured as the rate of loss of SWC per day, confirmed this difference in water consumption strategies. The water absorbed from soil appeared better retained in the (thick) leaves of *A. sagittata* than in the leaves of *A. nemorensis* (F_1, 195_ = 85.4682, *p* = 0.000139, **Fig-S5**). The two species, however, did not differ in stomata density and size (**Fig-S6**). Once wilted, plants were re-watered four days after the onset of wilting, and recovery was defined as the ability to form a new leaf within two weeks. *A. sagittata* showed a significantly higher recovery rate than *A. nemorensis*, with individuals being seven times more likely to form new leaves post-wilting. The proportion of recovered plants was significantly lower in *A. nemorensis* compared to *A. sagittata* (∼50% vs. ∼90%, F_1,197_ = 36.453, *p* = 1.08e-07, **Figure 1G**), highlighting the higher drought resilience of *A. sagittata*.

**Figure 1:**
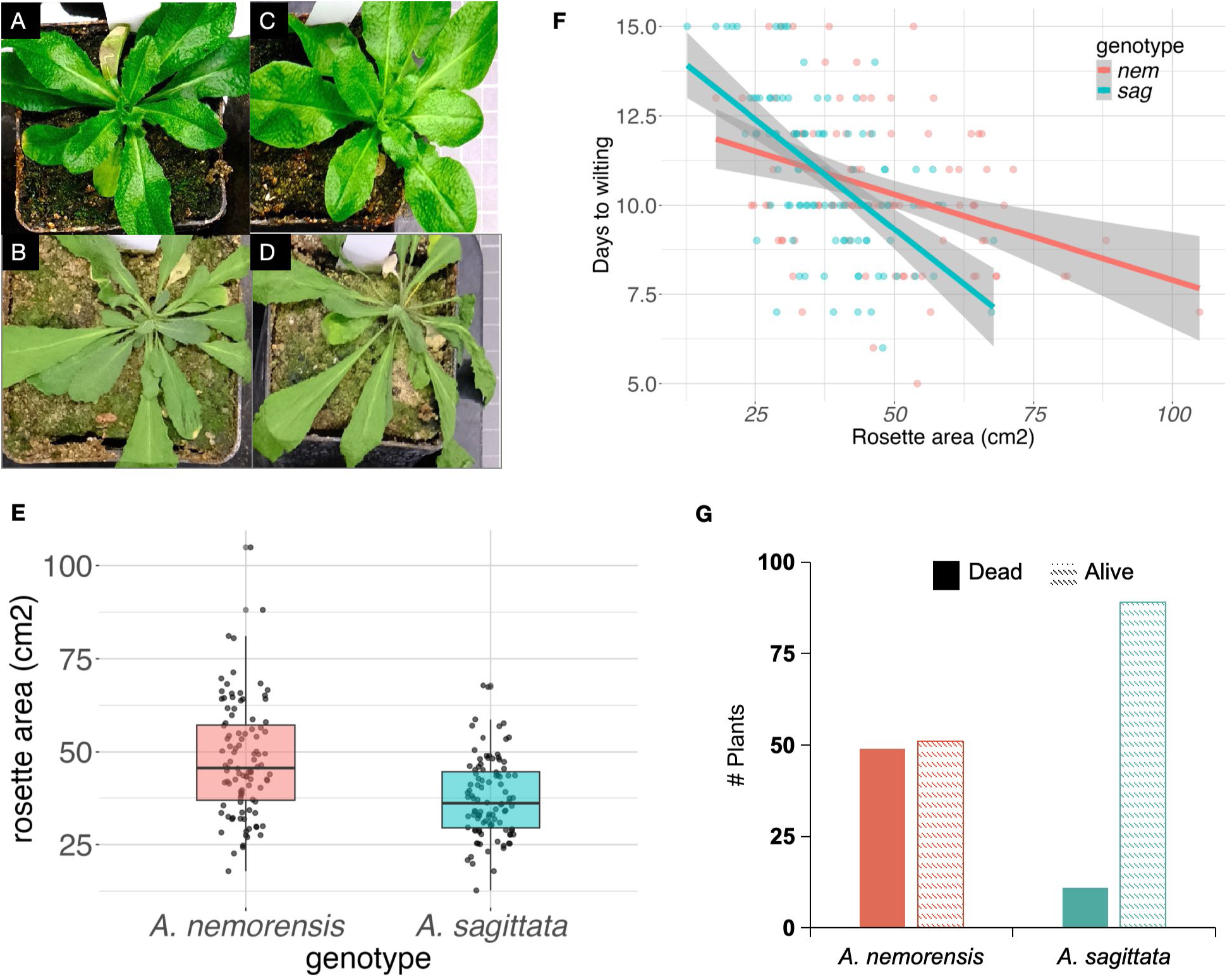
Phenotypes under well-watered conditions and during wilting in *Arabis sagittata* and *Arabis nemonresis*. *A. sagittata* (A, B) and *A. nemorensis* (C, D) plants before and after the drought treatment, respectively. (E) Boxplot shows that the rosette area of *A. nemorensis* is significantly larger than that of *A. sagittata* (*p* = 9.26e-08). (F) Days to wilting as a function of rosette area is explained by an interaction between rosette area and species (*p* = 0.00804). (G) Number of survivors after wilting and re-watering shows *A. sagittata* recovers at a higher rate (*p* = 1.08e-07) than *A. nemorensis*.

We also scored damage after recovery. In *A. sagittata*, more than half of the plants showed a degree of damage between 0 and 2 (0 = no damage, 6 = dead), whereas in *A. nemorensis*, more than half of the plants showed damage scores above 4 (F_1, 197_ = 27.767, *p* = 3.14e-16**, Fig-S4**). These results confirmed that *A. sagittata* tolerates soil dehydration and wilting better than *A. nemorensis*. In order to determine the genetic architecture of these strong differences, the two lines were used as parents to generate an interspecific segregating population, and 300 individuals of the F3 generation were genotyped using reduced-representation sequencing. When we followed the same protocol as above, about 75% of the F3 individuals survived. However, no significant QTL for survival was detected (**supplementary methods, Fig-S7**). These findings indicate that a complex polygenic basis underlies the difference in drought tolerance.

### Almost one-third of expressed genes respond differently to stress in the two species

To better understand the molecular changes underpinning differences in tolerance to drought, we quantified gene expression changes at the onset of wilting and after recovery. A principal component analysis indicated that transcriptome data clustered first by species and then by stress (**supplementary text, Fig-S8**). We used DESeq2 to identify genes with significant changes in expression for both species (**Fig-S9A-E**).

A total of 5,571 genes in *A. nemorensis* (adjP ≤ 0.05; fold-change ≥ 0.1) and 6,359 genes in *A. sagittata* (adjP ≤ 0.05; fold-change ≥ 0.1) were significantly differentially expressed during wilting versus control (**Table-S2 and Table-S3**). Similarly, when comparing recovery to control, 2,448 genes in *A. nemorensis* (adjP ≤ 0.05; fold-change ≥ 0.1) and 3,866 genes in *A. sagittata* (adjP ≤ 0.05; fold-change ≥ 0.1) had not returned to their pre-stress expression levels (**Table-S2 and Table-S3**). In total, 3,054 and 874 genes responded similarly in both species after wilting and after recovery, respectively **(Figure 2A and 2B).** Yet, for 3,980 genes (adjP ≤ 0.05; fold-change ≥ 0.1) both species differed in their expression response at wilting versus control, whereas 1,973 genes (adjP ≤ 0.05; fold-change ≥ 0.1) differed in their response to recovery versus control **(Table-S2 and Table-S3).**

**Figure 2:**
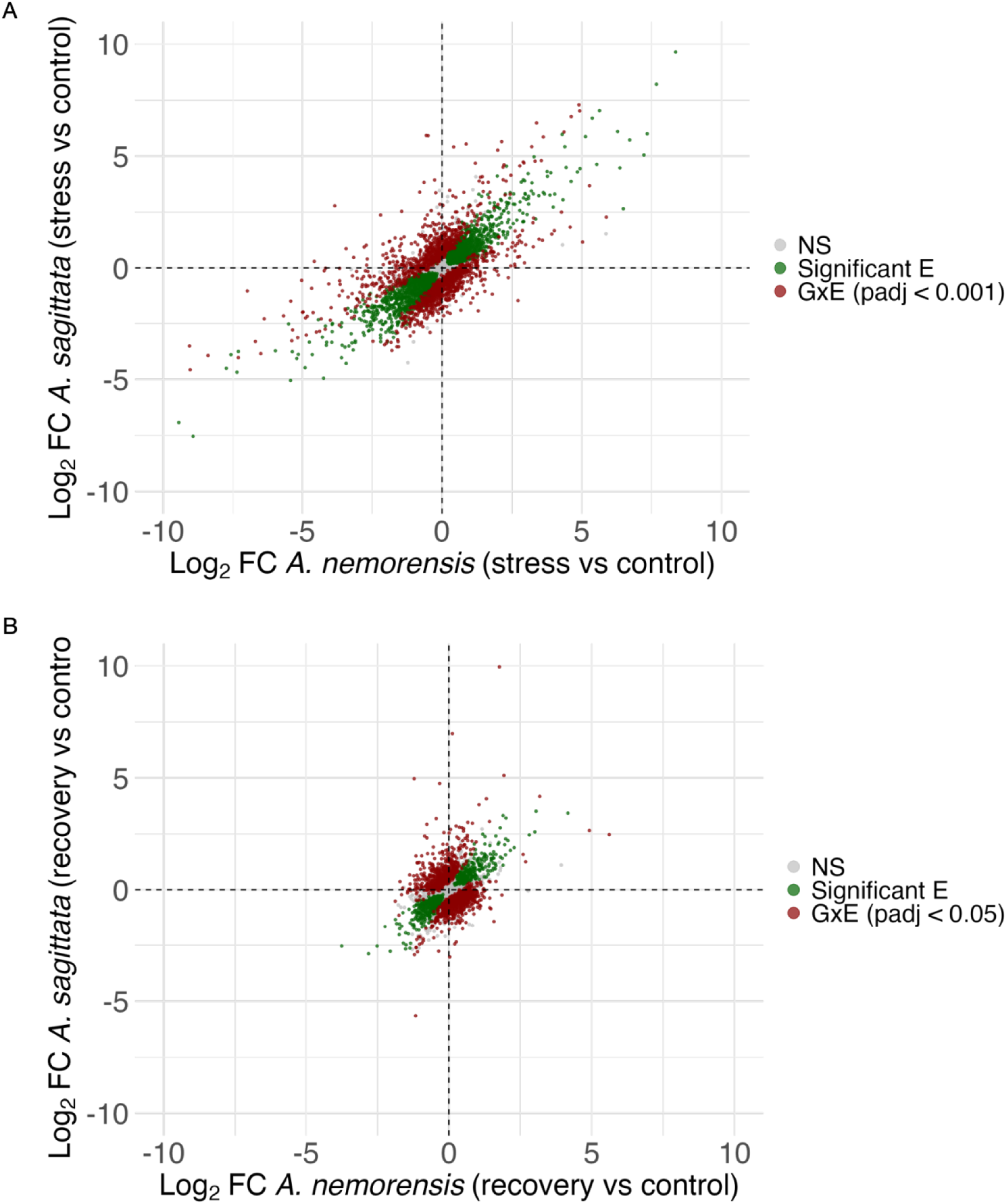
Byplot showing the stress response in *A. sagittata* as a function of the stress response in *A. nemorensis* (A) at the onset of wilting and (B) after recovery. Genes responding similarly are displayed on the diagonal (green), genes that respond differentially are shown in red. Genes that do not respond to stress in either species are shown in gray.

### Species activated distinct functions in responses to drought stress

Enrichment analysis in GO categories indicates that specific gene sets show enhanced responses in *A. sagittata* compared to *A. nemorensis,* (**Table-S4,** **Figure 2A**). Both species sensed and responded to the stress. Genes up-regulated during wilting in both species were enriched in functions related to response to stomatal movement (*p* = 0.0017), abscisic acid (*p* = 0.0034), and transcription by RNA polymerase II (*p* = 0.0034). Since *A. sagittata* performed better in response to stress than *A. nemorensis,* and recovered better, we used *A. nemorensis* as a reference in the following analysis and asked whether genes whose response differed from those of *A. nemorensis* were enriched in specific molecular functions and/or EPM clusters. We found that genes with increased expression in response to stress were strongly enriched in functions previously described as related to stress responses, yet the strongest functional enrichments were found among genes that were more downregulated during and after stress in *A. sagittata* compared to *A. nemorensis*, suggesting that *A. sagittata* survives in higher numbers than *A. nemorensis* by shutting down gene expression programs.

Among the genes upregulated at wilting in *A. nemorensis*, those that were upregulated at a higher level in *A. sagittata* compared to *A. nemorensis* were significantly enriched in several molecular functions related to stress, including genes responsible for the biosynthesis of alcohol (*p =* 0.00043), as well as genes that respond to light intensity (*p =* 0.00172), that respond to salt stress (*p =* 0.00189), that regulate cellular responses to hypoxia (*p* = 0.00202), and that respond to water deprivation (*p* = 0.00241). Conversely, the genes that responded less in *A. sagittata* compared to *A. nemorensis* were strongly enriched in functions such as the cellular response to red–far-red light (*p =* 0.00087), protein refolding (*p =* 0.00179), the metabolism of organic hydroxy compounds (*p* = 0.00382), the catabolism of small molecules (*p* = 0.00382), and the folding of chaperone-mediated proteins (*p =*0.0039). Among the genes that were upregulated in response to wilting in *A. nemorensis*, several responded in the opposite way in *A. sagittata* and were downregulated in response to stress. These genes were strongly enriched in functions related to the organization of chloroplasts (*p =* 2.70E-07), the initiation of translation (*p =* 4.40E-05), translation (*p =* 5.20E- 05), and the organization of thylakoid membranes (*p =* 0.00025).

Among the genes downregulated at wilting in *A. nemorensis*, those that were downregulated at a lower level in *A. sagittata* compared to *A. nemorensis* were strongly enriched in translation (*p =*3.00E-14), protein import into chloroplast stroma (*p =*5.30E-06), chloroplast organization (*p =* 1.20E-05), embryo development ending in seed dormancy (*p =* 0.00036), and the biosynthesis of hemes (*p =* 0.00039). Conversely, the genes that were less downregulated in *A. sagittata* compared to *A. nemorensis* were those involved in the regulation of transcription (*p =* 0.00062), the innate immune response (*p =* 0.00113), phosphorylation (*p* = 0.00192), the response to nematodes (*p* = 0.00371), and the biosynthesis of glucosinolates (*p* = 0.00371). Among the genes that were downregulated at wilting in *A. nemorensis*, several responded in the opposite way in *A. sagittata* and were upregulated. These genes were enriched in functions related to the response to salicylic acid (*p =* 1.60E-05), the response to molecules of bacterial origin (*p =* 0.00018), the cellular response to hypoxia (*p =* 0.0002), and the hormone-mediated signaling pathway (*p* = 0.0005).

After recovery, both species up-regulated genes related to the biosynthesis of flavonoids (*p* = 0.0022), the regulation of stress (*p* = 0.0027), the compound containing anthocyanin (*p* = 0.0038), and the pathway that mediates sugar signaling (*p* = 0.004). We followed the same logic as above to understand the functions that differentiated the responses of the two species (**Table-S5,** **Figure 2B**). Among the genes that showed an increased expression after recovery in *A. nemorensis*, those that responded more in *A. sagittata* compared to *A. nemorensis* were significantly enriched in the metabolism of starch (*p =* 7.00E-05), the response to oxygen-containing compounds (*p =* 0.00042), and the response to the metabolism of lipids (*p =* 0.00216) and amides (*p =* 0.00277). Conversely, those that were less upregulated in *A. sagittata* compared to *A. nemorensis* were significantly enriched in response to chemicals (*p =* 0.012), the metabolism of sulfur compounds (*p =* 0.013), and the response to cadmium ions (*p =* 0.014). Among the genes that were upregulated after recovery in *A. nemorensis*, several responded in the opposite way in *A. sagittata* and were downregulated after recovery. These genes were very strongly enriched in functions related to translation (*p =* 1E-30), cytoplasmic translation (*p =* 4.60E-10), chloroplast organization (*p =* 4.70E-06), ribosome assembly (*p* = 3.30E-05), and translational elongation (*p* = 3.30E-05). Thus, in the most drought-tolerant *A. sagittata*, not only were functions related to translation less upregulated at wilting, but they also appeared more suppressed at recovery compared to *A. nemorensis*.

Among the genes downregulated after recovery in *A. nemorensis*, those that were downregulated at an even lower level in *A. sagittata* as compared to *A. nemorensis* were significantly enriched in the metabolism of glucose (*p =* 5.20E-08), photosynthesis (*p =* 8.40E-08), and the biosynthesis of hexose (*p =* 5.20E-05) and chlorophyll (*p =* 9.10E-05). Conversely, the genes that were less downregulated in *A. sagittata* compared to *A. nemorensis* were enriched among those involved in the cellular responses to decreased oxygen levels (*p =* 0.0017), to oxygen levels (*p =* 0.0017), and to hypoxia (*p =* 0.0017). Among the genes that were downregulated after recovery in *A. nemorensis,* several responded in the opposite way in *A. sagittata* and were upregulated. These genes were enriched in functions related to responses to water deprivation (*p =* 0.0002), and to stomatal movement (*p =* 0.0047), and the metabolism of starch (*p =* 0.0014), Altogether, this analysis highlighted that during the drought stress *A. sagittata* strongly shuts down functions related to translation and chloroplast organization, a pattern that remains after recovery. On the other hand, alcohol biosynthesis and starch metabolism stood out as functions that were more strongly activated in *A. sagittata* compared to *A. nemorensis* upon wilting and after recovery, respectively.

### EPM content identifies functional targets of gene regulation in response to stress

Having detected markedly different transcriptome responses to stress in the two species, we sought to determine the molecular underpinning of these differences. After training models to predict gene expression (**Fig-S10a-b; Table-S6**), we used DeepCRE to identify expression predictive motives (EPMs) separately for each sampling time point and each species. In total, we identified 307 EPMs, in forward and reverse orientation (614) (**Table-S7**). Of these EPMs, 148 and 159 were extracted for *A. nemorensis* and *A. sagittata*, respectively. For *A. nemorensis,* we identified 54, 48, and 47 EPMs for control, wilting, and recovery conditions, respectively. For *A. sagittata*, we identified 58, 44, and 57 EPMs. The number of identified EPMs did not differ between species and conditions (χ2(2)=0.95, *p*=0.62). Based on sequence and position similarities (**Table-S7**), EPMs could be grouped into 25 clusters (**Table-S8**). EPM clusters can be shared across phylogenetically distant species such as *Arabidopsis thaliana* and *Zea mays* (Peleke et al., 2024). The EPM composition of the dataset reflects the diversity of sequence motives associated with active or repressive gene expression and thus the diversity of regulatory networks activated. Many EPM clusters were shared between *A. nemorensis* and *A. sagittata* and were found under all conditions, such as EPM cluster 13 (**Figure 3a-b**, **Table-S8**). EPMs in cluster 13, for example, predicted high transcript levels, resembled binding sites for GAGA-binding transcription factors like BPC1, and were preferentially positioned near transcription start and termination sites (TSS/TTS) **Table-S7**). Ubiquitous EPMs reflect regulatory motifs associated with modules that regulate basal expression; such modules are shared between species and active in response to both stress and control conditions.

**Figure 3:**
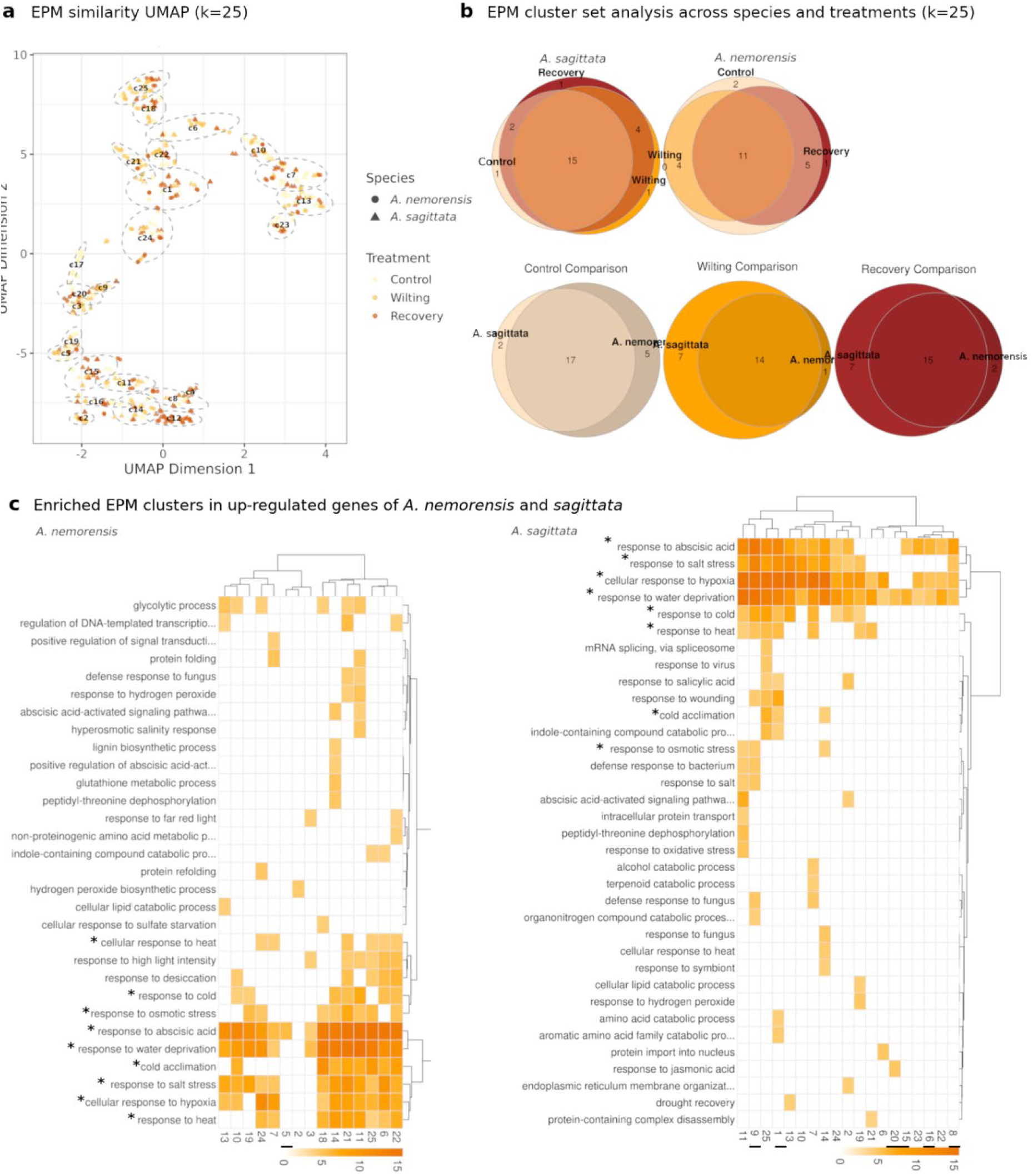
Expression Predictive Motifs (EPMs) clustered by similarity are linked to differentially expressed genes across Arabis species and treatments. (a) Importance scores, including gene and flanking regions, were aggregated into contribution weight matrices, which are referred to as expression predictive motifs (EPMs) using TF-MoDISco (Shrikumar et al., 2018). These EPMs were clustered into 25 groups, following similarity search by Pearson Correlation Coefficient (Peleke et al. 2024), and using k-means clustering, and visualized via UMAP representation. (b)We performed set analysis on EPM clusters to find those that were shared or unique to specific species and treatments. The Venn diagram shows a large overlap, indicating that most EPM clusters are shared among the different species and treatments. (c) Clustered heatmaps show EPM clusters among upregulated biological processes under wilting conditions of Arabis species. EPM clusters are significantly enriched (<0.1E-4 -log10 P of Fisher’s exact test) with gene ontology groups whose expression changes significantly during wilting. Shared gene ontology (GO) terms across the Arabis species are signified with an asterisk, while unique EPM clusters are signified by an underscore. While some EPM clusters are enriched in only one GO term, others are enriched in multiple GO terms.

The comparative analysis of EPM clusters detected in different species and at different stages in the drought-stress response can reveal whether species differ in the regulatory networks they activate during and after stress (**Table-S9)**. EPM cluster 5 was associated with genes with low transcript abundance in response to wilting in *A. nemorensis*. In this cluster, an EPM with 85% similarity to the KAN1 binding site, a repressor of auxin biosynthesis, was found in downstream intragenic and flanking extragenic regions, suggesting a transcriptional repression mechanism specific to drought stress in the species that is less likely to survive wilting **(Table-S7,** Huang et al., 2014). In addition, cluster 5 EPMs occurred in genes enriched in the biosynthesis of indoleacetic acids and glucosinolates, suggesting that the transcription factor binding to these EPMs represses the production of secondary metabolites (*p*= 4.20E-05 and 1.50E-06, respectively) (**Fig-S12**). EPMs with similarity to cluster 5 were absent from transcripts expressed in *A. sagittata*, and as mentioned above, glucosinolate-related genes were less downregulated by stress in this species, indicating a difference in how these genes are regulated.

In contrast to observations in *A. nemorensis,* gene expression in *A. sagittata* in response to wilting depended on a diverse set of EPM clusters (1, 8, 9, 15, 16, 20, 23, **Figure 3c, Table-S10**), none of which explained gene expression in *A. nemorensis* under the same conditions. *A. sagittata’s* response thus appears to be controlled by a larger number of regulators. Its response includes EPMs with a high similarity to experimentally characterized TFBS, such as activation via GATA8 sites (EPM cluster 1) in upstream intragenic regions, IDD family C2H2/C4 zinc finger sites (EPM cluster 16) near the TTS, DOF5.1 (EPM cluster 15), described as a family of C4 zinc finger types, or repression via ARF14 binding sites (EPM cluster 20) in upstream intragenic regions. Interestingly, EPM clusters 8, 15, and 23, all linked to high transcript levels in *A. sagittata* in response to wilting, did not show substantial similarity to known TFBSs, indicating that the drought response in *A. sagittata* may involve regulators that have not yet been discovered.

In order to identify candidate transcription factors associated with differential gene expression in response to wilting, we investigated EPMs enriched in genes that are regulated similarly (**Table-S11**). The low expression in *A. nemorensis* of stress-responsive genes containing EPMs of cluster 5 pointed to the high activity of a repressor with binding properties similar to MYS1, a MYB-SHAQKYF repressor of wax biosynthesis that was associated with high survival rates in response to wilting (Liu et al., 2022). Enrichment of EPMs of cluster 7 and 10 among genes that were more downregulated in response to stress in *A. sagittata* compared to *A. nemorensis* indicates strong activity of the transcription repressor HSFA6a, which exhibits 93% similarity to EPMs in this cluster. The repressor activity of HSFA6a has been associated with high rates of survival to drought (Hwang et al., 2014). EPMs with similarity with GATA, KAN1, Abi4, IDD5, DOF3.4, DOF5.8, or members of the ethylene response factor family point to additional candidate regulators that may explain the difference in gene regulation and contribute to high rates of drought survival in *A. sagittata* (**Table-S11**).

### *A. sagittata* drought response associates with expression variation of microRNA408

As one the most differentially expressed genes is a DEA(D/H)-box RNA helicase family protein gene, that is down-regulated in wilting only in *A. sagittata* (**Fig-S12)**. DEA(D/H)-box RNA helicases are known to regulate miRNA biogenesis and RNA splicing in *Arabidopsis* and are involved in stress responsive gene regulation (Guan et al., 2013, Xu et al. 2023). Since transcriptional response to wilting and other stresses is also linked to miRNA regulation (Jeong et al. 2013; Singh et al. 2023) we further hypothesized that miRNAs may play a key role in differential tuning gene expression between *Arabis* species.

We sequenced short RNA of samples from the same drought and recovery treatments and anlayzed variation in miRNA expression levels with DESeq2. We detected 20 expressed miRNAs in *A. nemorensis* and/or *A. sagittata*, of which 16 very low expression (**Table-S13).** Three of the highly expressed miRNAs did not differ significantly in expression between conditions and species. Only miR408 showed significant interspecific difference in expression in response to stress and after recovery (interaction b/w condition ANOVA *p-value* = 0.00114), being significantly upregulated in *A. sagittata* in drought conditions, but not in *A.nemorensis*. This miRNA has a well-documented role in drought tolerance in *A. thaliana* (Ma et al., 2015).

The miR408 genes of both *Arabis* species displayed sequence differences supporting the notion that they differ in cis-regulation. A 6.2 kb insertion 453 bp upstream of the mature MIR408 (about 340 bp upstream of the transcription start site) in *A. nemorensis* (**Figure 4A-C**). BLAST analysis against the NCBI database showed that this insertion represents an unknown low copy number non-LTR retrotransposon. Additionally EPMs from clusters 10 and 14 mapped directly onto the miR408-3p and -5p regions within the precursor locus, respectively, being candidates for differential expression levels. However, we identified three key variations altering the EPM contents of the two alleles in the gene flanking regions (**Figure 4A, and D-F, Table-S12**). At position -426, a single nucleotide polymorphisms (SNP) distinguished the *A. nemorensis* (T) and *A. sagittata* (C) alleles. The *A. nemorensis,* variant created a motif matching EPMs associated with low transcript abundance (**Figure 4D**). In contrast, the *A. sagittata* variant created a motif matching EPMs associated with high transcript abundance (**Figure 4B**). The EPMs of the *A. sagittata* alleles resembled binding sites of UBP3 (epm_Asag_SW_p0m20 cluster 11, ∼82% similarity) and IDD family members (epm_Asag_SW_p0m22 cluster 16, ∼85% similarity) (**File-S1; Table-S7**). The third EPM (epm_Anem_SW_p1m05, cluster 11, ∼93%), only matching the *A. nemorensis* variant does display similarity to transcription factor binding sites of CDF5 of the DOF family in the JASPAR database (Rauluseviciute et al. 2024). Prediction of TFBS using PCBase 2.0 derived from CHiPseq and DAPseq data in *A. thaliana* (Chow et al. 2024), however, predicts that the T/C mismatch between *A. nemorensis* and *sagittata* affects binding of SUPPRESSOR OF PHYB-4#3 (87% similarity score), respectively (**File-S1, Table-S12**). Therefore the two alleles differed in predicted affinity to several transcription factors.

**Figure 4:**
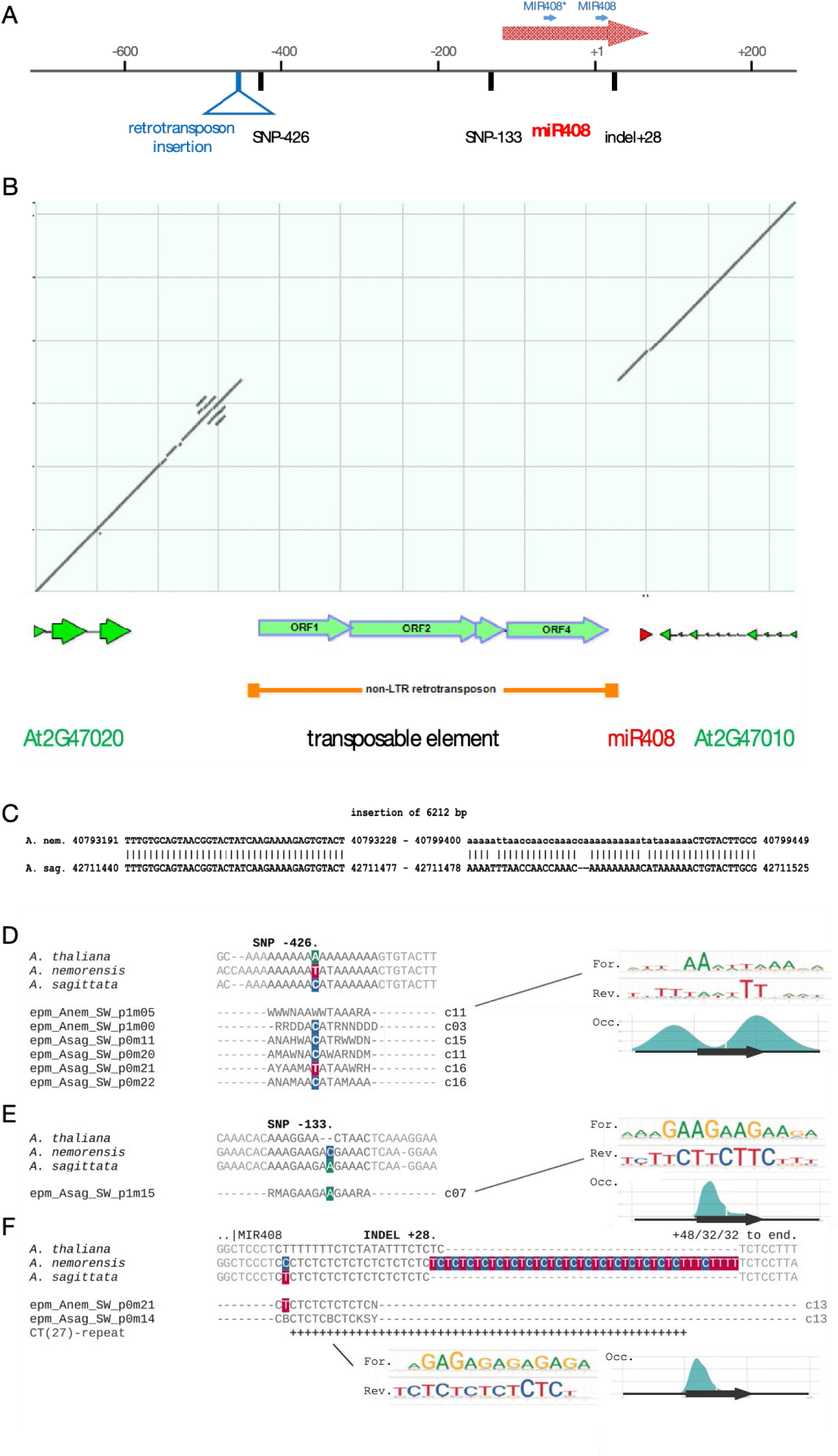
Transposable element insertion in chromosome 4 upstream of the *Arabis nemorensis* miR408 gene. (A) Schmematic representation of sequence alterations between *Arabis nemorensis* and *A. sagittata* in the miR408 region. (B) Dot plot of sequence alignment of the genomic regions encompassing the orthologues of the *Arabidopsis thaliana* genes At2G47020, At2G47015 (miR408), and At2G47010 showing a 6212 bp insertion 453 bp upstream of the mature miR408 (about 340 bp upstream of the miR408 transcription start site). The schematic representation under the dot plot indicates the locations and orientation of the genes. (C) Sequence comparison around the retrotransposon insertion site. Numbers are positions on the respective *A. nemorensis* and *A. sagittata* chromosomes. (D) A SNP located 428 bp upstream of miR408 influences the mapping of EPMs identified in wilting treatments. The EPM epm_Anem_SW_p1m05, with shown motif in forward and reverse (For./Rev.) orientation and preferred occurrence (Occ.) relative to gene region (black arrow), associated with reduced transcript levels. This is a candidate proposed to explain the observed differential gene expression of miRNA408. (E) A SNP positioned 133 bp upstream of miR408 impacts the mapping of epm_Asag_SW_p0m15, exclusively detected in *A. sagittata* during wilting treatment. This EPM is associated with elevated transcript abundance and predominantly found in intragenic upstream regions. (F) An INDEL mutation situated 28 bp downstream of the miR408 start site and 36 bp upstream of the end of miR408 precursor mRNA, engenders an extended 27 fold CT-repeat motif in *A. nemorensis*, aligning with EPM cluster 13. An example of this EPM is epm_Anem_SW_p0m21, which correlates with high transcript abundance and exhibits a positional preference for intragenic upstream regions.

A second upstream region of the miRNA408 gene start, features a SNP at position -133 (*A. nemorensis*: C; *A. sagittata*: A). This variant created a motif in *A. sagittata* that matches an EPM of cluster 7 with high similarity to TFBS for heat shock factor HSFA6A (93%) and NAC transcription factor-like 8 (87%) (**Figure 4E**, **Table-S12**). PCBbase 2.0 confirmed this result by predicting the presence of TFBS with varying similarity to HSFA1A in this region for *A. nemorensis,* (91%) and *A. sagittata* (95%) (Chow et al. 2024). The palindromic consensus sequence AGAA--TTCT is known to be bound by HSFA1A and HSFA6A. The *A. sagittata* sequence shows greater similarity to this consensus, even though it has mismatches at positions 7 (A) and 8 (A).

Finally, we also detected an INDEL mutation within the hairpin region. According to predicted secondary structures of miRNA408 (Kozomara et al. 2019), a CT-hexamer region, which the *Arabis* species share with *Arabidopsis thaliana*, is involved in complementary binding of the folded RNA. In *A. nemorensis,* the CT-nucleotide has been duplicated compared to *A. sagittata* (**Figure 4F**). Consequently, the INDEL mutation and extension of the CT-polymer in *A. nemorensis,* may also affect RNA secondary structure of miRNA408.

### miR408 targets stress response genes that respond differently in the two species

Using TargetFinder (Bo and Wang 2005), a tool for predicting miR408-targets among the differentially expressed genes. We identified nine genes were potentially targeted by miRNA in *Arabis sagittata* and sixteen genes in *Arabis nemorensis,* respectively (**Fig-S13**). Only in *A. sagittata*, the significantly expressed miR408 potentially targeted stress genes. Two of the genes that miR408 potentially regulates, At5G13650 and At5G65280 are down-regulated in response to stress in *A. sagittata*, the species in which miR408 is activated, but not in *A. nemorensis* (**Fig-S14**). At5G13650 (SUPPRESSOR OF VARIEGATION 3) is involved in the regulation of response to oxidative stress (Baxter et al., 2007) and At5G65280 (GCR2-LIKE 1) is a gene of unknown function (Guo et al., 2008). But inconsistent expression patterns were also observed for other putative targets of miR408. We also noted that the DeepCRE model outputs did not identify EPMs similar to the target sequence of miR408. The number of known or predicted miRNA targets may be too low to train the model on this regulatory layer.

## Discussion

Plant species growing in grasslands offer unique opportunities to study drought tolerance, as they have evolved to balance the biotic demands imposed by competition within dense communities with the challenges of abiotic stress. A rare example of sympatry between *A. nemorensis*, a species confined to floodplain meadows, with its close relative from dry calcareous grasslands, *A. sagittata*, offered excellent models for studying drought tolerance in competitive species. We found that both *A. nemorensis* and *A. sagittata* are particularly drought-tolerant because they do not wilt until the SWC drops below 5-10%. In comparison, the model plant *Arabidopsis thaliana* wilts at 10-17% SWC but does not recover; its relatives, *A. halleri* and *A. lyrata,* wilt at 18-20% SWC (Bechtold et al., 2016; Bouzid et al., 2019). At those levels, *A. sagittata* and *A. nemorensis* show no sign of wilting. Given known trade-offs between growth and stress tolerance, this ability to withstand such extreme soil drought is remarkable for two species that are highly competitive in dense meadows. (Donath et al., 2006). However, though both species wilted at similar SWC levels, *A. sagittata*, which prefers dry calcareous meadows, recovered markedly better than *A. nemorensis.* Because no quantitative trait locus could be identified, the molecular basis of this difference is thought to be polygenic, as is common in complex traits (Sella and Barton, 2019).

Determining the genetic basis of polygenic ecological traits is always a challenge and more so in non-model species. To gain insights into the molecular basis of these differences, we monitored changes in the transcriptome and miRNA expression at the onset of wilting and after recovery, and compared the reactions of the two species. Functional enrichments among differentially expressed genes indicate species have an array of molecular and physiological responses to maintain homeostasis and adaptive function under stress (Zhu et al., 2016; Nakashima and Shinozaki, 2013; Shaar-Moshe et al., 2017). Using a machine-learning approach that identifies sequence features predicting high or low expression in each species and at each stress stage, we observed that the predicting features differ between species and stress conditions (Peleke et al., 2024). These differences indicate that the two species activate not only common reactions but also different regulatory networks. Sequences of genes that differ in how they react in the two species highlight families of regulators, all of which have known roles in the responses of plants to abiotic stress. These regulators likely form the basis of the polygenic underpinnings of the improved drought tolerance of *A. sagittata*.

*A. sagittata* mounts a stronger response for several classes of drought-related genes such as those involved in response to salt stress, hypoxia, or water deprivation as well as for those involved in the biosynthesis of alcohol. The exogenous addition of ethanol on *A. thaliana* plants has been shown to increase tolerance to drought, presumably by triggering neoglucogenesis and the accumulation of sugars, glucosinolates, and amino acids (Bashir et al., 2022). Since hypoxia favors the production of alcohol in plants, it is tempting to speculate that these biochemical reactions contribute to its improved survival rates. In addition, the rice gene OsFAR1, which is involved in the biosynthesis of fatty alcohols, promotes drought in rice and decreased cuticular permeability in *A. thaliana* (Guan et al., 2023). We further observed that sequences similar to the binding site of the repressor MYS1 were enriched among genes that were expressed at high levels in *A. sagittata*. Recently MYS1 has been associated with the diurnal regulation of wax biosynthesis in *A. thaliana* (Liu et al., 2022). Since *A. sagittata* has thick leaves that are less prone to water loss during dry-down, further work could also test whether differences in MYS1 activity underpin its ability to withstand drought.

*A. sagittata* also differed from *A. nemorensis* in its sharply decreased transcription of genes involved in the organization of chloroplasts. This pattern of expression may reflect the ability of drought-tolerant species to maintain photosynthesis and thus allocate sugars to other organs. Alternatively, replenishing starch reserves helps reinitiate growth and keep energy balance, and *A. sagittata* displayed significantly less damage in response to stress (Thalmann and Santelia, 2017). After recovery, *A. sagittata* maintained a high expression of genes enriched in functions related to the metabolism of starch as well as to the response to lipids, oxygen-containing compounds, stomatal movement, osmotic adjustment, and developmental programming (functions that were all downregulated in *A. nemorensis* in response to stress). The analysis of expression predictive motifs further identified several GATA transcription factors via the enrichment of cognate sequence motifs in *A. nemorensis*. Since this family of transcription factors regulates photosynthesis and development (Schwechheimer et al., 2022), variation in these factors may underpin the strong differences in how chloroplasts and genes regulate photosynthesis that were observed in *A. sagittata* during wilting.

Beyond the regulation of gene function in response to stress, the role of miRNA-mediated regulation during drought stress has been increasingly recognized as crucial (Filipowicz et al., 2008). Our findings revealed that miR408 was the only miRNA that is differentially expressed between the species, with *A. sagittata* sharply upregulating miR408 during wilting (unlike *A. nemorensis*, which did not modify its expression). The putative targets of miR408 included genes whose orthologs in *A. thaliana* are known to regulate reactive oxygen species (ROS) (Balyan et al., 2017; Yang et al., 2024) and in abscisic acid (ABA) signaling (Yang et al., 2024). MiR408 is a highly conserved miRNA across plant species (Pan et al., 2018) and has a demonstrated role in stress mitigation, particularly in rice (Zhou et al., 2010; Yao et al., 2022) and *Arabidopsis thaliana* (Ma et al., 2015). It has also been implicated in regulating responses to nutrient stress, such as the assimilation of sulfur assimilation and the adaptation to arsenic in Arabidopsis (Kumar et al., 2023). The alleles of *A. sagittata* and *A. nemorensis* had marked differences in cis-regulation. In addition, the *A. nemorensis* allele had a large insertion upstream from the transcript, and the *A. sagittata* allele harbored EPMs matching binding sites of the heat shock regulator HSFEA, which suggests potential cis-regulatory differences between alleles. Yet, the absence of a QTL for drought tolerance in this region suggests that the differential regulation of miRNA408 is caused by species differences in the polygenic regulatory network activated in response to stress and not only by these putative cis-regulatory elements.

The two genotypes used for this study were collected in the same floodplain meadow near the Rhine River in Riedstadt, Germany, a location where *A. sagittata* and *A. nemorensis,* form natural hybrids (Dittberner et al., 2019). These genotypes may not represent the entirety of the species-wide diversity of *A. sagittata* and *A. nemorensis*, but they do indicate species differences observed in a single location where intermixing occurs (Dittberner et al., 2022). Drought in floodplain grasslands, which experience fluctuating cycles of flooding and drought, is increasing in intensity due to climate change and river management (Barendrecht et al., 2024). In this study, we document the markedly higher drought tolerance of *A. sagittata* compared to *A. nemorensis*. In conclusion, the polygenic differences shown by these two genotypes’ responses to drought are likely to contribute to shape the outcome of interspecific hybridization in this system (Dittberner et al., 2022; Rahnamae et al., 2025). Future studies will be needed to explore whether improved drought tolerance evolved at the expense of competitive ability (Grimes et al., 1977; Mount et al., 2023).

## Acknowledgments

We thank K. Bell for help during the experiment, DAAD for funding, as well as DFG for funding with grant number ME2741/13-1, Project-ID 456082119 – TRR 341 and EXC-2048/1 project ID 390686111. Emily Wheeler for editorial assistance.

## Author contributions

ASK performed the experiment and data analysis. JDM designed experiments, supervised data analysis and interpretation, and edited the manuscript, with input from GS and IC. SMZ generated bioinformatic pipeline for the characterization expression predictive motifs across species with consultation of JJS, TA, NR, and LA helped in building the reference genome, LH instructed on the steps in QTL analysis, RW reviewed orthologue identification. ASK and JDM wrote the manuscript, with input from all authors.

## Data availability

The raw RNA-seq data of the transcriptome and small RNA-seq used in this study have been submitted to the ENA website with the accession number PRJEB78710.

## Supplementary data

Supplementary text (methods, figures, and tables) Other supplementary tables and code can be accessed via github respository: https://github.com/Abdubidopsis/Arabis_drought_transcriptome

