## Supplementary information for "The complex molecular basis of enhanced stress resilience in extreme drought-tolerant *Arabis* grassland species"

### Supplementary Methods

#### Growth conditions

Eight weeks after harvest, seeds were stratified on wet paper for five days at 4°C in darkness (Dittberner et al. 2019). A total of 100 germinated seedlings per species were then transplanted individually into 7x7x8 cm (about 3.15 in) pots each filled with 360g of a well-homogenized mixture of VM soil (Einheitserde VM, Einheitserde Werkverband eV, Germany; 60-70 % peat and 30-40% clay), perlite and ceramics (clay granules). Pots were split in ten blocks (trays) and distributed across three shelves of a CLF growth chamber (Perkin Elmer, USA), with 14h light at 20°C, 10h dark at 16°C, 100 µmol $m^{-2}s^{-1}$ light intensity supplemented with 10 min dark-red light at the end of the day: Trays were rotated across the shelves throughout the experiment.

#### Sampling for RNA Extraction

For RNA extraction, young leaves (close to the center of rosette) were sampled from 50 plants at the end of the acclimation period (control sample), at the appearance of wilting symptoms (wilting sample), and after recovery. For the wilting time point, plants were sampled as they showed signs of imminent wilting. Leaf material was collected at the same hour of the day (four hours Zeitgeber time).

#### Detailed protocol of phenotypic measurements

Rosette area was quantified on day zero of water withdrawal using the open-source software ImageJ and its Rosette Tracker plugin, originally designed to measure *Arabidopsis thaliana* growth by counting pixels and converting them into $mm^2$ (De Vylder et al., 2012). Initial leaf thickness and leaf thickness at wilting were measured on two medium-sized leaves per plant. Each leaf was marked with ink to ensure that the same leaf was measured at both time points. Leaf lamina thickness was assessed using a digital ruler (HOLEX, Hoffmann Group, Knoxville, TN, USA) with an accuracy of ±0.02 mm. SWC was monitored daily until wilting, as described in the main text. The rate of water loss was calculated as the rate of SWC decay from day zero of the dry-down experiment until wilting. Recovery was scored two weeks after wilting. Recovery time was determined by counting the number of days between re-watering and the emergence of a new fresh leaf. Additionally, plant images taken at the beginning of water withdrawal and during the recovery phase were used to quantify damage severity. Damage was visually assessed using a six-level scale reflecting the percentage of lost leaf area, changes in leaf color, and leaf damage or senescence, where 1 indicated minimal damage and 6 represented plant death. Stomatal density and stomatal length were measured using an optical microscope on approximately five plants per species. For each plant, three leaves were selected, and three spots per leaf were analyzed. Stomatal traits were quantified at the end of the acclimation period following the protocol described by Paccard et al. (2014). Leaf area was measured using ImageJ software on three medium-sized leaves per plant for ten plants per species.

**mRNA and miRNA extraction protocol**

Leaf tissue was homogenized using a Precellys Evolution homogenizer (Bertin Technologies, Montigny-Le-Bretonneux, France) for 3 × 10 seconds at 6800 rpm, with intermittent cooling in liquid nitrogen to prevent thawing. RNA was extracted using the Macherey-Nagel Plant RNA extraction kit (Macherey & Nagel, Düren, Germany), and RNA concentration was measured with a NanoDrop 2000c spectrophotometer (Thermo Scientific, Waltham, MA USA). RNA quality and quantity were assessed using an Agilent 2100 Bioanalyzer (Agilent Technologies, Palo Alto, CA, USA) with RNA Nano chips. Only high-quality RNA samples (OD260/280 = 1.8~2.2, OD260/230 ≥ 2.0, RIN ≥ 6.5, 28S:18S ≥ 1.0, >50 μg) were retained for sequencing. All mRNA samples were sequenced in a single batch using the Illumina HiSeq 4000 platform at Azenta (Genewiz) Leipzig, Germany, following the manufacturer's protocol.

For small RNA sequencing, leaf samples were collected following the protocol described above. Small RNA was extracted using the Qiagen RNeasy Plant Mini Kit (Qiagen, Hilden, Germany). RNA concentration was initially assessed using the NanoDrop 2000c, while RNA quality and quantity were evaluated using the Agilent Tape Station (Agilent Technologies, Palo Alto, CA, USA). Only high-quality RNA samples (OD260/280 = 1.8–2.2, OD260/230 ≥ 1.6, RIN ≥ 6, 28S:18S ≥ 1.0, >50 μg) were selected for library preparation. A total of 24 leaf RNA samples were sequenced using Illumina SE50 technology at the Cologne Center for Genomics (CCG), Germany.

**Bioinformatics analysis of RNA Transcriptome**

For transcriptome data processing, we first used the FastX-toolkit from the FastQC package (v0.11.4) for raw sequence quality assessment, trimming, and filtering, following the approach described by He et al. (2016). Low-quality nucleotides were removed from the 3' ends of the sequences using a Phred score threshold of 20 (t = 20) and a minimum read length of 50 bp. Sequences were reverse complemented using the fastx reverse complement function to ensure consistent trimming at both ends. Reads with >90% of bases below the quality threshold and paired end reads missing one valid pair were discarded from further analysis.

We used Hisat2 to map the trimmed and filtered reads to the *Arabis nemonresis* high quality reference genome with reference in the European Nucleotide Archive (ENA) (Project id: PRJEB89863) (Rahnamae et al. 2025). The transcriptome sequencing yielded an average of 20 million paired-end reads per sample with a read length of 150 bp. Read quality was assessed using Samtools (version 1.3.1), applying the command "samtools view -q 10" to retain high-quality, uniquely mapped reads with a correct mapping probability of ≥ 90%, thereby keeping on average 86% of the reads. RNA integrity was verified using a custom R script to confirm uniform transcript coverage and ensure that RNA degradation did not bias expression estimates. Gene expression quantification was performed using HTSeq-count, and DESeq2 (Bioconductor version: Release 3.5) was used to identify differentially expressed genes (DEGs) between conditions (Love et al., 2014). We applied the Wald test to compute ps, using the model: ~ species + timepoint + species:timepoint, where the factor species had two levels (*A. nemorensis,*, and *A. sagittata*), and the factor timepoint included three conditions: (1) leaves sampled at 60% soil moisture, (2) at 5% soil moisture, and (3) after recovery. Genes were considered significantly differentially expressed if they met the thresholds of adjusted p (≤ 0.05) and log2-fold change (≤ -0.1 or ≥ 0.1). Contrasts were applied to identify DEGs in both *A. nemorensis*, and *A. sagittata* across all three conditions. Scripts can be found in GitHub repository: https://github.com/Abdubidopsis/Arabis_drought_transcriptome

**Small RNA sequencing analysis**

Small RNA sequencing yielded 20 million single end reads per sample with a read length of 50bp. We used Bowtie to map the trimmed and filtered reads to the *A. nemorensis* reference genome. We used the samtools (version 1.3.1) "samtools view -q 10" to select the unique reads with highly quality with a probability of correct mapping of 90%. We filtered out the longer reads (>30bp) and kept small RNA reads for further analysis. On average 90% of small RNA reads were successfully mapped to the reference genome. HTSeq-count was used to measure the read counts for small RNA. The DESeq2 Bioconductor package from R (Bioconductor version: Release 3.5) was used to find the position of all small RNAs as well as 21nt and 24nt small RNA on the PCA for control, wilting and recovery.

For miRNA target prediction, we used TargetFinder to identify putative miRNA targets in *A. nemorensis* and *A. sagittata* under stress conditions. First, a list of known plant miRNAs was downloaded in FASTA format from miRBase (https://www.mirbase.org/). Small RNA reads were mapped to these known miRNAs and subsequently remapped to the reference genome, allowing the identification of known miRNAs and their potential target genes. Low-quality miRNAs were filtered out based on the TargetFinder score function, retaining only miRNAs with a score ≥ 4. To explore the relationship between miRNAs and differentially expressed genes (DEGs), I overlapped putative miRNA targets with DEGs identified in transcriptome samples collected in control, at the onset of wilting, and after recovery. The overlap was visualized using a Venn diagram generated online (http://bioinformatics.psb.ugent.be/) and tested for enrichment in differentially expressed genes using a hypergeometric test. Additionally, miRNA expression and its association with DEGs were visualized in R, while the Cytoscape platform (https://cytoscape.org/) was used to generate interaction networks between miRNAs and their target genes.

**Identification of expression predictive motifs (EPM)**

Deep learning models for the prediction of transcript levels were trained for *Arabis nemonresis* and *Arabis sagittata* RNAseq experiments control, wilting and recovery, respectively. We used the strategy for training laid out by Peleke and colleagues and trained species and treatment specific single-species reference models (SSR) and shuffled-sequence controls (SSC) for control (Peleke et al. 2024). As defined in (Peleke et al. 2024), the cis-regulatory sequences are considered to be the 1000 bp upstream of the gene's transcription start site (TSS), the 1000 nt downstream of the gene's transcription termination site (TTS), 500 bp downstream from the transcription start site (TSS), and 500 bp upstream from the transcription termination site (TTS). The extracted sequences from both ends are one-hot encoded, separated by a pad (pad size is 20), and concatenated. Across species and treatments, models were trained in a binary classification task on low and high transcript profiles. To ensure robust classification of the two classes across different experimental conditions, the classification strategy was adjusted compared to Peleke et al 2024. Here transcript levels were labelled as "high" transcript profiles if these belonged to the upper 20th percentile of the base-10 log-Median of transcript per million (log10MedTPM). The same number of genes was sampled from a pool of genes with a log10MedTPM of zero and gene ranked that were minimally larger than zero (1:1). Accordingly, the classes were balanced and percentile threshold were more similar across different species and treatments (log10MedTPM 20th percentile thresholds for *A. nemorensis* and *A. sagittata* control, wilting and recovery, respectively: 1.288, 1.233, 1.296, 1.246, 1.267, 1.338). In addition to the work of Peleke et al 2024, the performance of models was evaluated by loss, accuracy, auROC and auPR (**Table-S6**).

Area under receiver operator characteristic (auROC) values above 0.8 show that information in the DNA sequence was identified that allows to effectively distinguish high and low expressed genes. For *A. nemorensis*, the best model was obtained after training on Chromosome 7 for control and wilting and on chromosome 2 for recovery. AuROC values reached 0.827, 0.845 and 0.854 for each model, respectively. For *A. sagittata*, the model trained on Chromosome 7 achieved best performance for control, whereas for wilting and recovery, the best models were obtained after training on Chromosome 5, with auROC reaching 0.813, 0.82 and 0.818, respectively (**Fig-S10a**).

### GO term enrichment analysis

*A. thaliana* orthologs of the annotated genes in *A. nemorensis* or *A. sagittata* were identified via Orthofinder, and genes lacking orthologs were excluded from this analysis (Emms and Kelly 2019). The comprehensive set of unique orthologs expressed in our dataset served as the background universe for Gene Ontology (GO) enrichment analysis. GO enrichment analysis was conducted utilizing the topGO (2.59.0) package in R (Alexa, Rahnenführer and Lengauer 2006). Emphasis was placed on the biological process ontology, employing the 'elim' algorithm alongside Fisher's exact test to account for GO hierarchy, utilizing org.At.tair.db (3.21.0) for annotations.

GO terms with a Fisher *p* below 0.0001 were deemed statistically significant (**Table-S10,** following He et al. 2016).

Functional enrichment analysis of DEGs was conducted independently for each species across the following comparisons: (1) wilting vs. control (upregulated genes), (2) wilting vs. control (downregulated genes), (3) recovery vs. control (upregulated genes), and (4) recovery vs. control (downregulated genes). Genes were further categorized based on their difference in fold-change response to stress in both species both at wilting and after recovery.

Since genes sharing EPMs are predicted to be co-regulated, we also used GO enrichment analyses to gain further insight into the molecular functions that were under the control of a common EPM. We focused on the EPMs extracted from the response to wilting and mapped their occurrence in the genome using BLAMM (Fostier 2020) and filtered using criteria analogous to those in Peleke et al., 2024 (BLAMM score >10) onto the annotated genomes of *A. nemorensis,* and *A. sagittata* (Rahnamae et al. 2025). EPMs exhibiting positional preference within 1000 nt upstream and downstream regions adjacent to the gene, or within the initial or terminal 500 nt of the transcribed regions, were assigned to the respective gene and filtered. Concurrently, selected DEG exhibiting up- or down-regulation, in conjunction with assigned EPMs of *A. nemorensis,* and *A. sagittata*.

EPM based clustering of DEGs provided an additional layer of functional grouping, relating shared gene functions and their regulation. Analyses were conducted to determine statistically significant GO terms associated with each EPM cluster, and clusters were subsequently ranked according to the magnitude of their GO enrichment signals, as determined by the count and mean negative base-10 logarithm of ps for significant terms. Subsequently, clustered heatmaps were generated using the pheatmap package (1.012) to visualize the enrichment significance, expressed as the negative base-10 logarithm of ps of GO terms across the distinct EPMs clusters for both up-regulated and down-regulated gene sets. The methodology assessed whether genes harboring sequences of a specific EPM cluster exhibited a higher frequency of annotation with a particular GO term than would be expected by chance, given the prevalence of that GO term within the background gene set. Consequently, a GO term was deemed statistically significantly characteristic of the gene set in which a specific EPM of a given cluster was observed (**Table-S10)**.

**Supplementary figures**

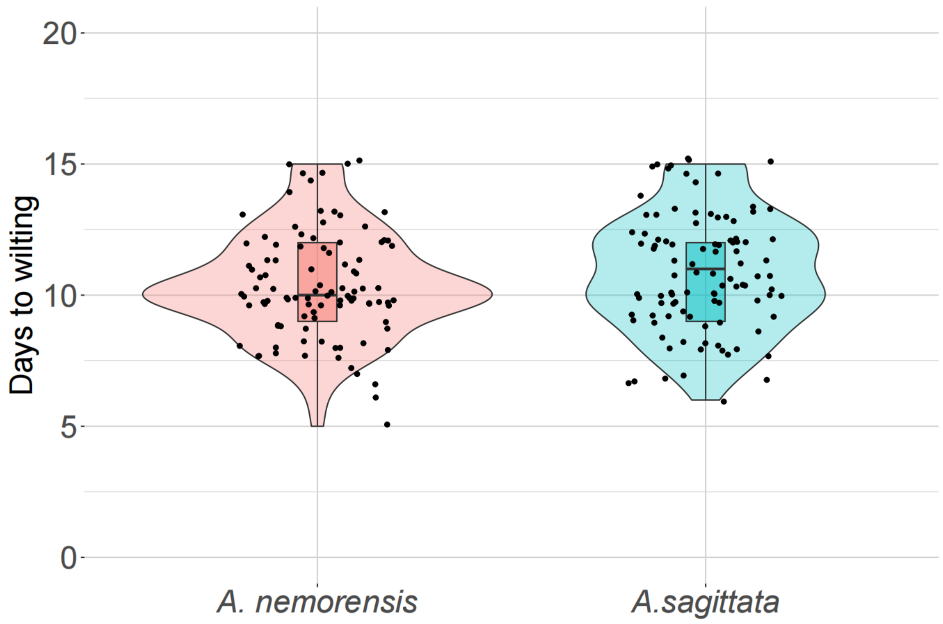

**Fig-S1:** Number of days a specie(s) required to wilt during the dry down experiment does not differ between species ($F_{1,197} = 2.5736$, $p = 0.239$).

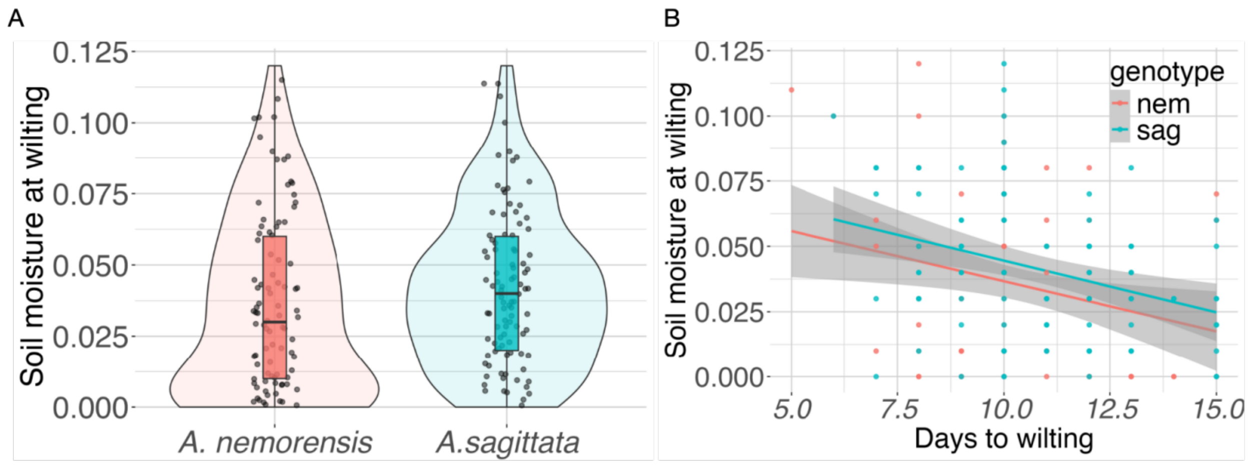

**Fig-S2:** Soil moisture when (A) the plants are about to wilt during the dry-down experiment does not differ between species, (B) Significant interaction was observed between SWC and days to wilting ($F_{1,197} = 21.608$, $p = 1.25e\text{-}12$).

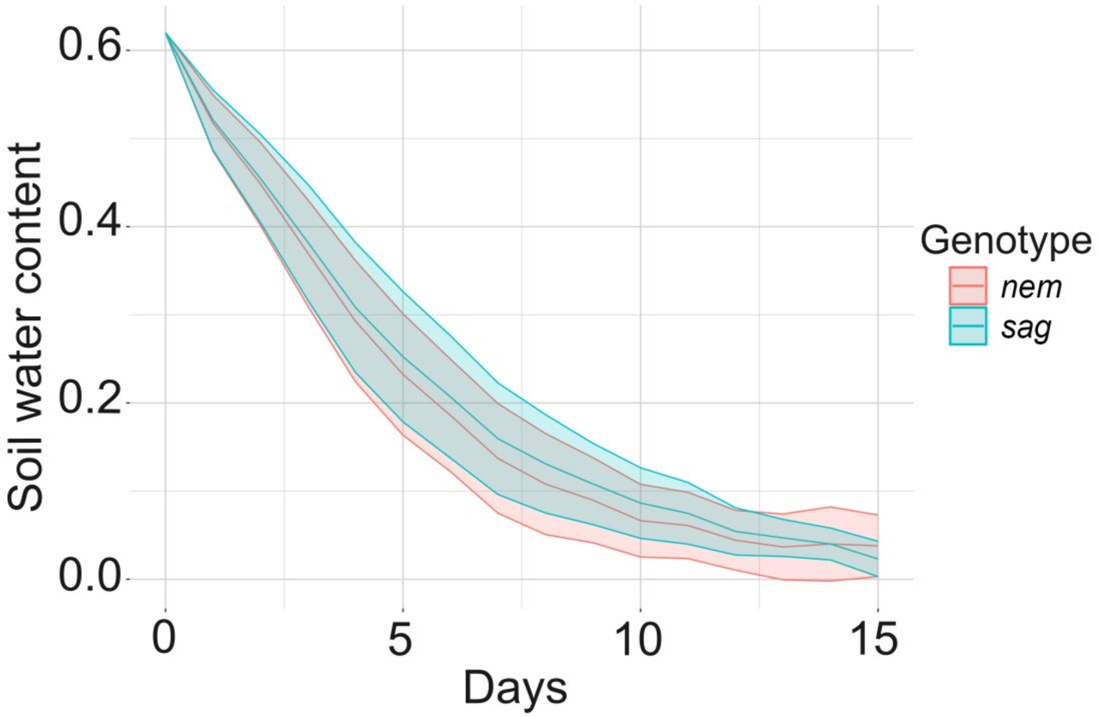

**Fig-S3:** Moisture loss per day during the dry down experiment for *Arabis sagittata* and *Arabis nemorensis* does not differ between species ($F_{1,\,1364.2}$= 0.4998, $p$ = 0.4799).

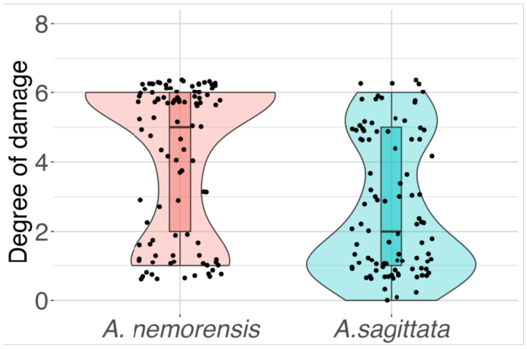

**Fig-S4:** Degree of damage (DoD) scored after recovery from wilting in *A. nemorensis* and *A. sagittata*. A general linear model was used to identify significant differences ($F_{1,\,197}$ = 27.767, $p$ = 3.14e-16).

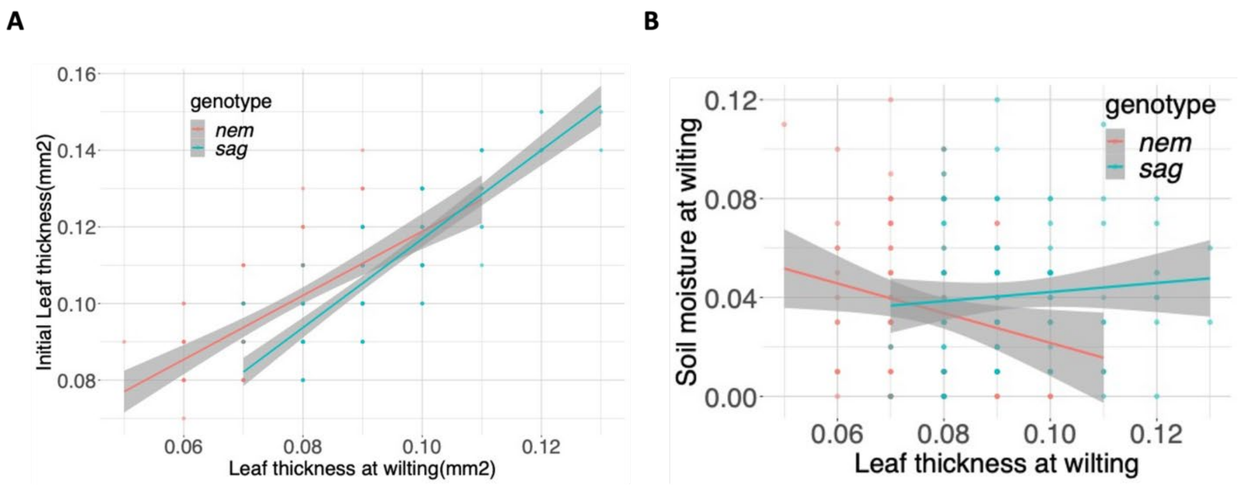

**Fig-S5: Variation in leaf thickness suggests better water retention in *A. sagittata***

(A) Variation in leaf thickness can be used to quantify variation in leaf water content. Initial leaf thickness and leaf thickness at wilting were more strongly correlated in *A. sagittata* plants compared to *A. nemorensis,* ($F_{1, 396} = 10.248$, $p = 0.00148$). (B) There was also a significant interaction between species and soil moisture at wilting on variation in leaf thickness at wilting ($F_{1, 195} = 85.4682$, $p = 0.000139$), which suggests that in *A. sagittata* the water absorbed from soil is more retained in the leaves than in *A. nemorensis*. The lines represent a linear regression smoothing where the shaded ribbons represent the standard error. Pink: *A. nemorensis,*, “nem”; Blue: *A. sagittata*, “sag”.

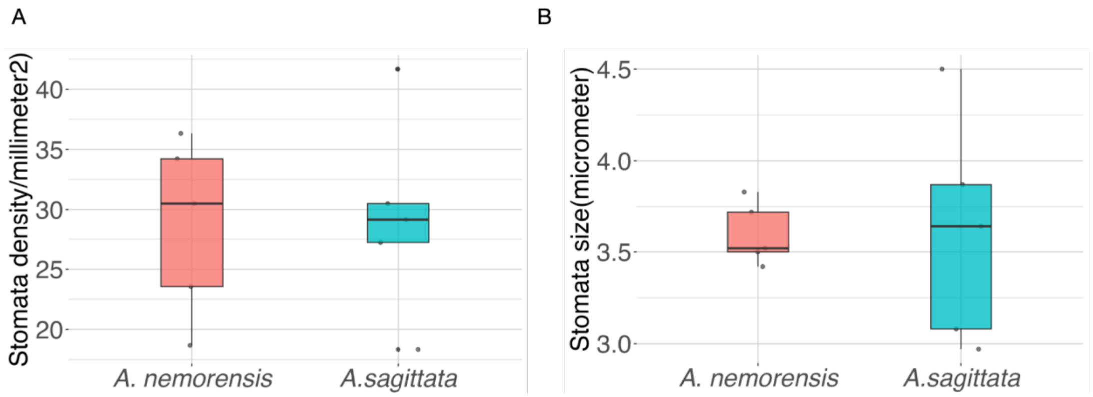

**Fig-S6: Species do not differ in stomata density and length**

We examined whether the two species exhibited constitutive differences in leaf stomatal patterning. Our analysis revealed no significant differences in stomatal density or stomatal size on the abaxial leaf surface. The average stomatal density was 29.38 stomata/mm² in *A. nemorensis,* and 28.66 stomata/mm² in *A. sagittata*, with no significant difference between species ($F_{1,5}$ = 0.0539, $p$ = 0.2213, **A**). Similarly, stomatal size did not significantly differ between species ($F_{1.5}$ = 0.0066, $p$ = 0.6431, **B**). Plot shows the (A) stomata density per millimeter square (B) Stomata size in micrometer in the leaves of *Arabis nemonresis,* and *Arabis sagittata*.

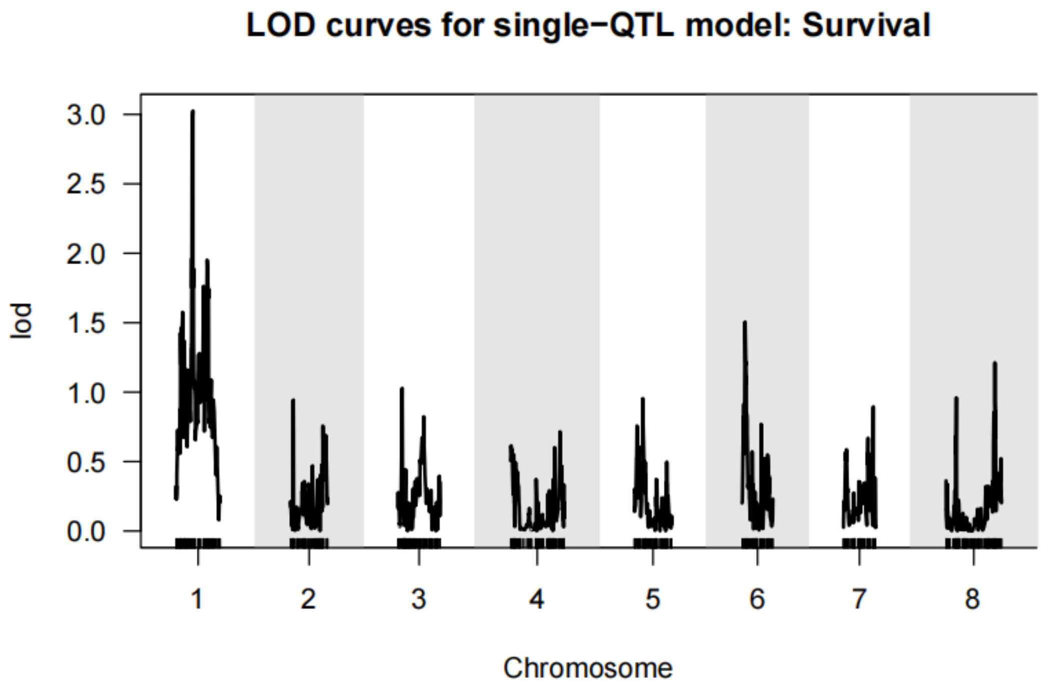

**Fig-S7: No significant QTL was detected for survival to drought**

Since both species differed in survival to drought, we conducted QTL mapping for survival. We grew an F3 population of 203 individuals for which a genetic map spanning eight chromosomes was constructed following previously described methods (Rahnamae et al. 2025). Individual plants were submitted to the dry-down treatment described above and survival was scored. Using a single-QTL model, we performed genome-wide scans with 5000 permutation tests, which established a significance threshold of LOD = 3.7 at a genome-wide significance level of $\alpha = 0.05$. No genomic region exceeded this threshold, indicating the absence of a major-effect QTL for survival in this population which also suggests the trait is polygenic. Figure shows the LOD score profiles for survival under drought stress across the genome. The x-axis represents chromosomes 1 through 8, while the y-axis shows the LOD score.

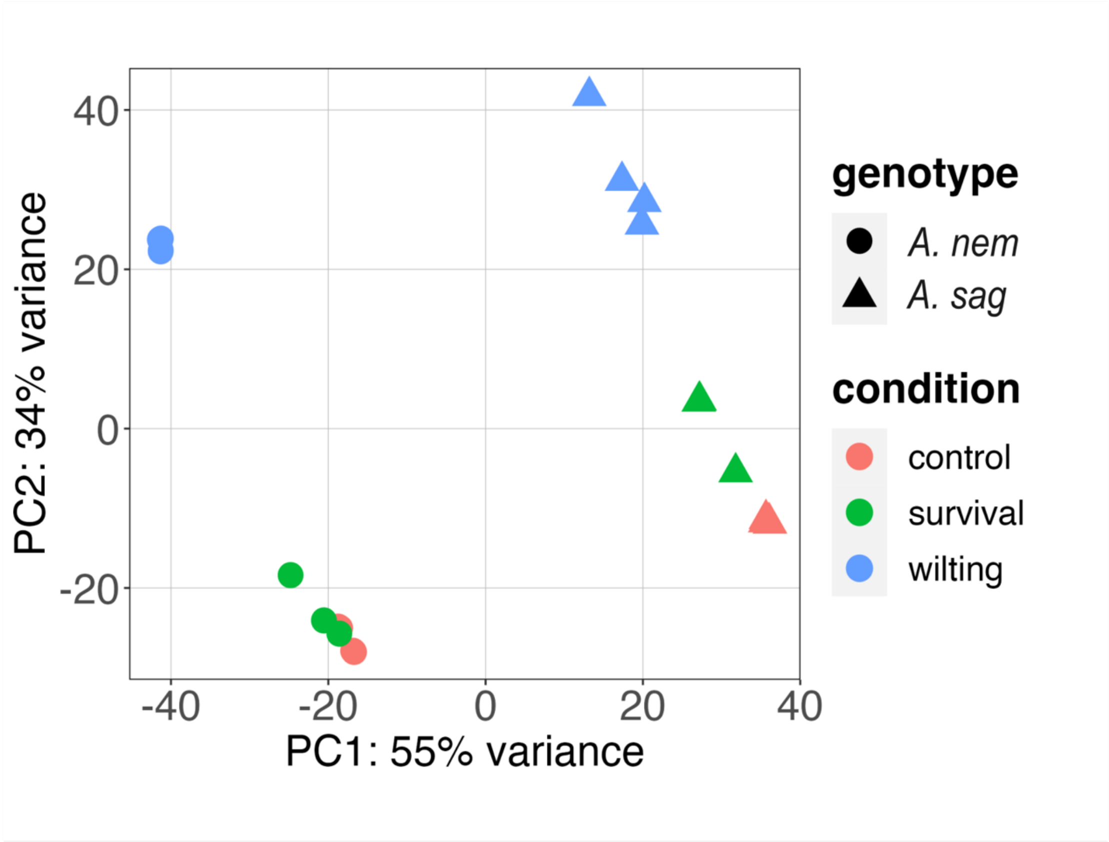

**Fig-S8: Transcriptome variance first due to species differences and then to stress conditions at the time of sampling**

We performed a principal component analysis (PCA) on variation in standardized read counts. Species clustered separately along the first principal component (55% variance), whereas the response to wilting separated samples along the second principle component (34% variance). Interestingly, samples collected after recovery did not appear to differ from control samples collected in well-watered conditions.

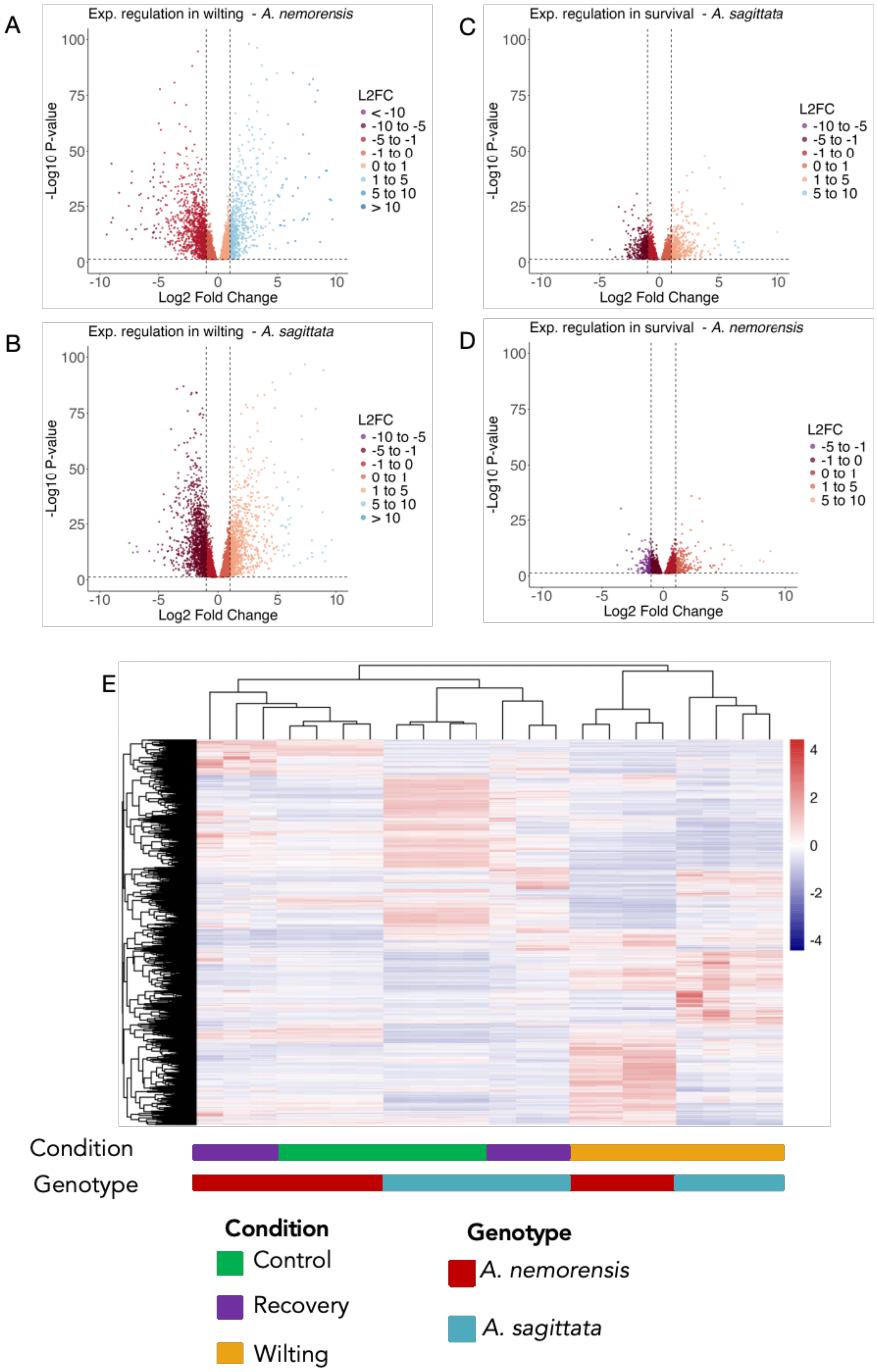
A
Exp. regulation in wilting - A. nemorensis
-Log10 P-value
Log2 Fold Change
L2FC
< -10
-10 to -5
-5 to -1
-1 to 0
0 to 1
1 to 5
5 to 10
> 10
B
Exp. regulation in wilting - A. sagittata
C
Exp. regulation in survival - A. sagittata
D
Exp. regulation in survival - A. nemorensis
E
Condition
Genotype
Control
Recovery
Wilting
A. nemorensis
A. sagittata

231

**Fig-S9:** Expression pattern of genes with a significant interaction between species and conditions of stress at the time of sampling (Likelihood Ratio test – See suppl. Method)**.** (A) *A. nemorensis,* at 5% vs 60% soil moisture, (B) *A. sagittata* at 5% vs 60% soil moisture, (C) *A. nemorensis,* at recovery vs 60% soil moisture, (D) *A. sagittata* at recovery vs 60% soil moisture, (E) The heatmap shows genes that respond differently to the conditions of stress between the two species.

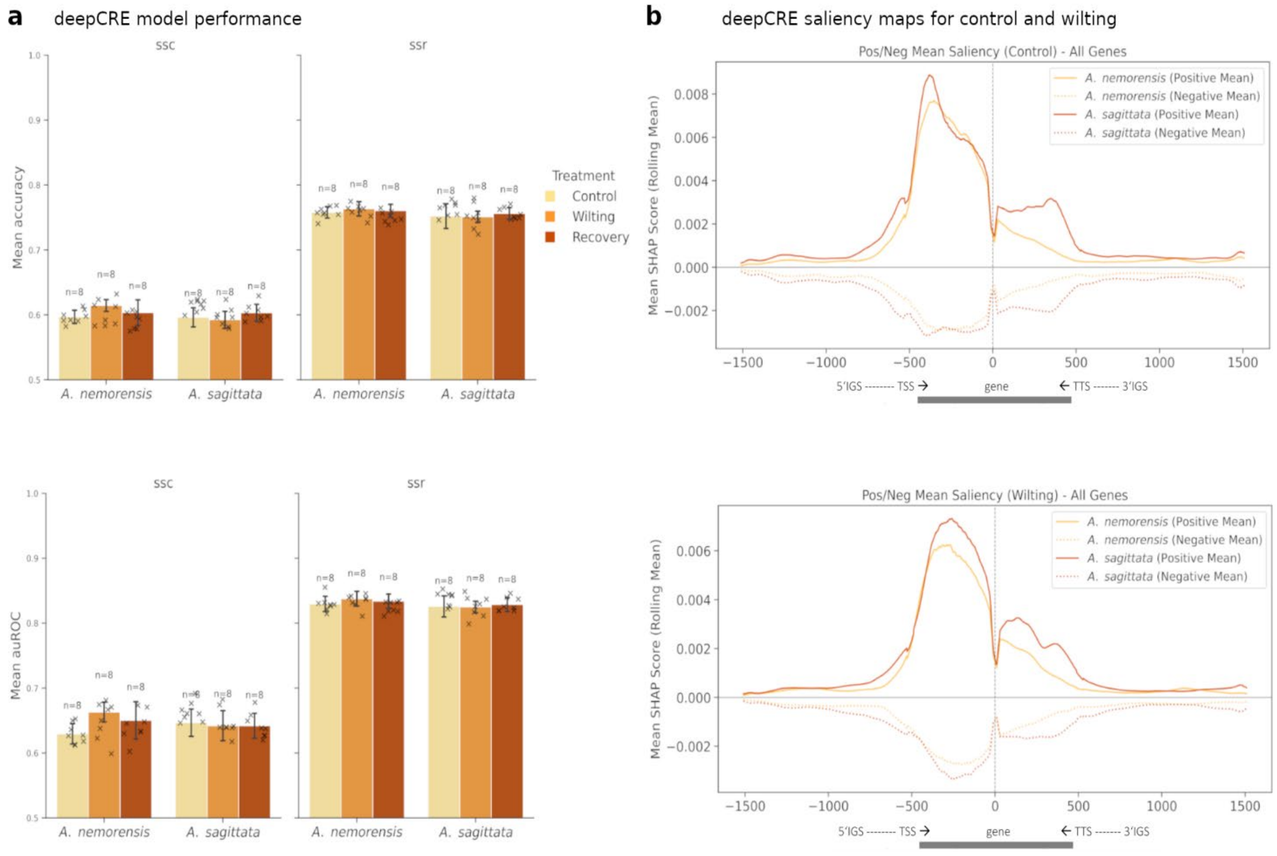

**Fig-S10:** Modeling gene transcript levels with high performance and feature extraction to identify predictive regulatory elements responsive to environmental conditions.

a) DeepCRE model evaluation of single species reference (SSR) and single nucleotide shuffle controls (SSC) demonstrates high accuracy and area under the Receiver Operating Characteristic (auROC) for *A. nemorensis* and *A. sagittata* across different environmental treatments: control, wilting, and recovery (for each chromosome n = 8). Individual data points are indicated by crosses within the barplots.

b) Saliency maps highlight important upstream and downstream regions around the genes transcription start and termination site (TSS, TTS) contributing to accurate model predictions using SHAPley (Lundberg and Lee 2017). These maps, generated for *A. nemorensis* and *A.*

*sagittata* across the control and wilting treatment, were used to compute averaged nucleotide importance per normalized position and expression predictive motifs (EPMs). These scores were calculated using a 40 base-pair (bp) rolling window. Positive SHAP scores indicate DNA regions that are important for predicting high gene expression, while negative SHAP scores show areas crucial for predicting low gene expression. Our analysis revealed that the average SHAP scores were consistently important for their respective predictions of high and low gene expression, and this pattern held true across all species and treatments studied.

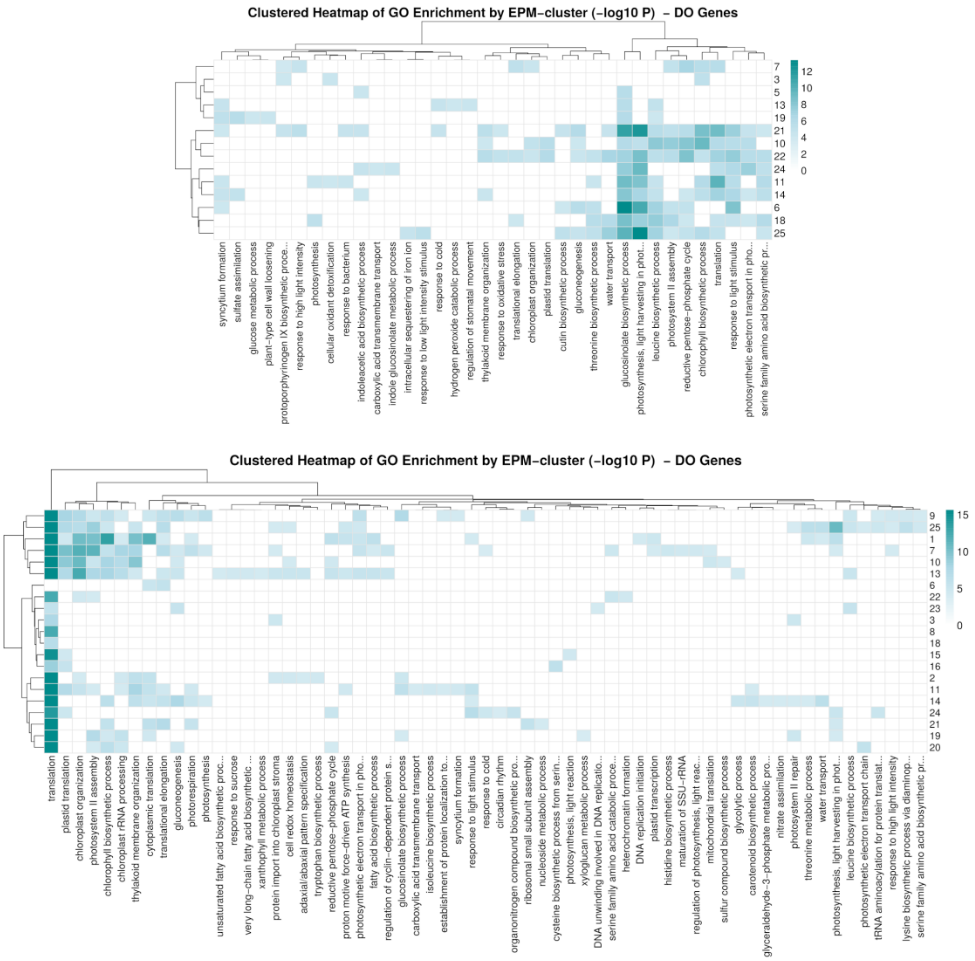

**Fig-S11:** Heatmaps for EPM cluster enrichment among down regulated biological processes of *A. nemorensis* and *A. sagittata* in response to wilting

Clustered heatmaps show EPM clusters among downregulated biological processes under wilting conditions of *Arabis nemorensis* (top) and *sagittata* (below). EPM clusters are significantly enriched (<0.1E-4 -log10 P of Fisher's exact test) with gene ontology groups whose expression changes significantly during wilting. While some EPM clusters are enriched in only one GO term, others are enriched in multiple.

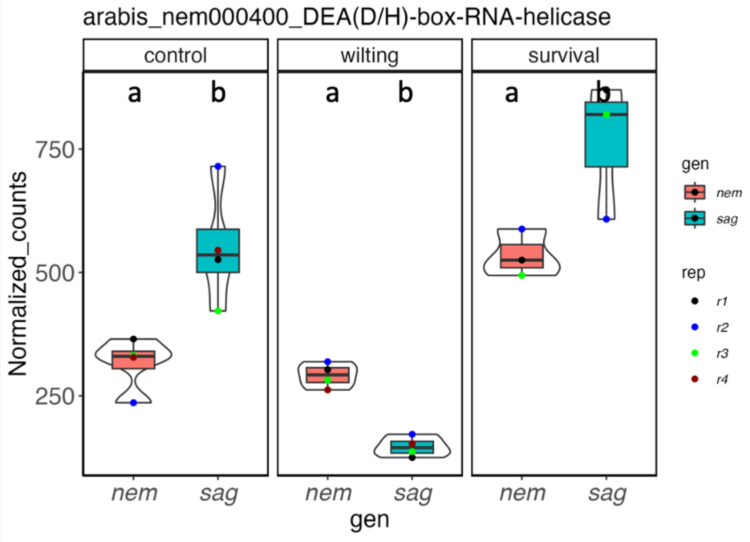

**Fig-S12:** Differential expression of the miRNA related Dea(D/H) box gene between two species in different conditions. The Arabidopsis thaliana ortholog encodes an RNA helicase with a role in miRNA biogenesis and RNA splicing (Xu et al. 2023).

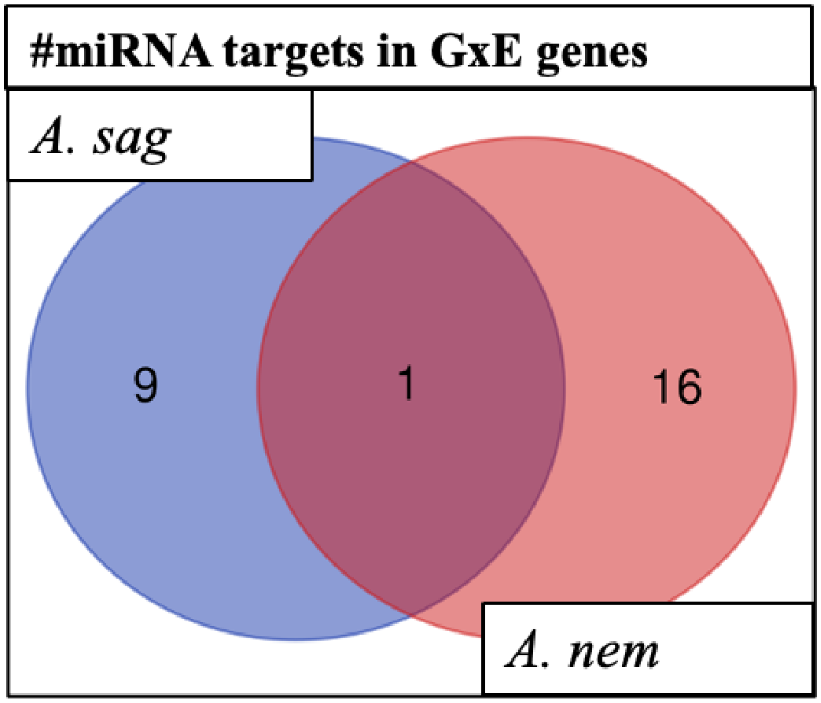

**Fig-S13: Overlaps in potential targets by expressed microRNAs in the two species.** Venn diagram shows number of miRNAs potential targets in GxE gene shows. We identified nine genes in stress were potentially targeted miRNA in *Arabis sagittata*, sixteen in A. nemorensis, and one gene potential target was found in both species. We then checked the expression of detected 20 miRNAs, 4 of which were expressed, while one miRNA (miR408) was differentially expressed and significantly expressed in A. sagittata. All of those miRNAs for . *nemorensis,* and *A. sagittata*, respectively were confirmed with database miRBase. Only in *A. sagittata* the significantly expressed miR408 targeted stress genes. While in *A. nemorensis* the genes were targeted by non-expressed or non-significantly expressed miRNA.

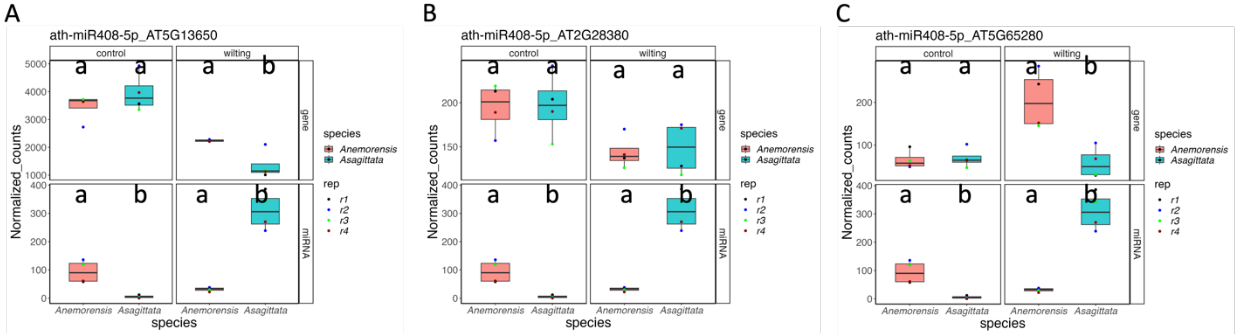

**Fig-S14:** Boxplot shows expression regulation of miR408 and potential targets (A) AT5G13650, (B) AT2G28380 and (C) AT5G65280.

### Supplementary tables

**Table-S1:** Phenotypes of *Arabis nemonresis* and *Arabis sagittata* measured during the dry down experiment. https://github.com/Abdubidopsis/Arabis_drought_transcriptome

**Table-S2:** Numbers of significantly differentially expressed genes of *A. nemorensis,* and *A. sagittata* during the dry-down experiment at contrasts of 5% vs 60% soil moisture and recovery vs 60% soil moisture.

| Condition | Specie | # genes Up | # genes Down |
|---|---|---|---|
| 5% vs 60% | *A. nemorensis* | 2825 | 2746 |
| | *A. sagittata* | 3236 | 3123 |
| Recovery vs 60% | *A. nemorensis* | 1371 | 1077 |
| | *A. sagittata* | 1925 | 1941 |
| GxE 5% vs. 60% | | 2009 | 1971 |
| GxE Recovery vs 60% | | 855 | 1118 |

**Table-S3:** Analysis of differential gene expression in *Arabis nemonresis* and *Arabis sagittata* in stress and recovery. https://github.com/Abdubidopsis/Arabis_drought_transcriptome

**Table-S4:** Genes responding to incoming wilting differently in *A. sagittata* compared to *A. nemorensis* at FDR<0.05 are enriched in several biological functional GO categories. Given are GO.ID: gene ontology identifier number, Term: description, number of annotated genes in this category, number of genes reacting in the respective way, expected number of genes, *p*-value of the enrichment. Enrichment were computed separately depending on the pattern of differential expression.

**Functions enriched among genes that respond identically to wilting in *A. sagittata* and *A. nemorensis***

| GO.ID | Term | Annotated | Significant | Expected | KS |
|---|---|---|---|---|---|

| | | | | | |
|---|---|---|---|---|---|
| GO:0010119 | regulation of stomatal movement | 31 | 19 | 10.56 | 0.0017 |
| GO:0009737 | response to abscisic acid | 170 | 79 | 57.89 | 0.0034 |
| GO:0006366 | transcription by RNA polymerase II | 113 | 57 | 38.48 | 0.0034 |
| GO:0045292 | mRNA cis splicing, via spliceosome | 10 | 8 | 3.41 | 0.0039 |
| GO:1905037 | autophagosome organization | 22 | 14 | 7.49 | 0.0042 |
| GO:0006814 | sodium ion transport | 5 | 5 | 1.7 | 0.0045 |
| GO:0090332 | stomatal closure | 16 | 11 | 5.45 | 0.0048 |
| GO:0048584 | positive regulation of response to stimu... | 69 | 34 | 23.5 | 0.0059 |
| GO:0006357 | regulation of transcription by RNA polym... | 89 | 42 | 30.31 | 0.0063 |
| GO:0048519 | negative regulation of biological proces... | 254 | 109 | 86.49 | 0.0074 |
| GO:0009395 | phospholipid catabolic process | 7 | 6 | 2.38 | 0.0077 |
| GO:0010305 | leaf vascular tissue pattern formation | 7 | 6 | 2.38 | 0.0077 |
| GO:0009788 | negative regulation of abscisic acid-act... | 17 | 11 | 5.79 | 0.0094 |
| GO:0009624 | response to nematode | 15 | 10 | 5.11 | 0.0099 |
| GO:0090333 | regulation of stomatal closure | 11 | 8 | 3.75 | 0.0101 |

**Functions enriched among up-regulated genes that are more up-regulated in *A. sagittata* than *A. nemorensis* at wilting**

| GO.ID | Term | Annotated | Significant | Expected | KS |
|---|---|---|---|---|---|
| GO:0046165 | alcohol biosynthetic process | 17 | 6 | 1.09 | 0.00043 |
| GO:0009642 | response to light intensity | 45 | 9 | 2.89 | 0.00172 |
| GO:0009651 | response to salt stress | 146 | 19 | 9.37 | 0.00189 |
| GO:0071456 | cellular response to hypoxia | 46 | 9 | 2.95 | 0.00202 |
| GO:0009414 | response to water deprivation | 127 | 17 | 8.15 | 0.00241 |
| GO:0045017 | glycerolipid biosynthetic process | 32 | 7 | 2.05 | 0.0033 |
| GO:0042548 | regulation of photosynthesis, light reaction | 11 | 4 | 0.71 | 0.00371 |

| | | | | | |
|---|---|---|---|---|---|
| GO:0006720 | isoprenoid metabolic process | 42 | 8 | 2.69 | 0.00426 |
| GO:0016567 | protein ubiquitination | 125 | 16 | 8.02 | 0.00508 |
| GO:0033993 | response to lipid | 236 | 25 | 15.14 | 0.00679 |
| GO:0000422 | autophagy of mitochondrion | 14 | 4 | 0.9 | 0.00968 |
| GO:0048584 | positive regulation of response to stimulent | 69 | 10 | 4.43 | 0.01097 |
| GO:0008299 | isoprenoid biosynthetic process | 31 | 6 | 1.99 | 0.01198 |
| GO:0046890 | regulation of lipid biosynthetic process | 15 | 4 | 0.96 | 0.01255 |
| GO:0006644 | phospholipid metabolic process | 61 | 9 | 3.91 | 0.01389 |
| GO:0006721 | terpenoid metabolic process | 32 | 6 | 2.05 | 0.01399 |
| GO:0009739 | response to gibberellin | 34 | 6 | 2.18 | 0.01868 |
| GO:0009644 | response to high light intensity | 25 | 5 | 1.6 | 0.01869 |
| GO:0019760 | glucosinolate metabolic process | 10 | 3 | 0.64 | 0.02197 |
| GO:0016143 | S-glycoside metabolic process | 10 | 3 | 0.64 | 0.02197 |
| GO:1902644 | tertiary alcohol metabolic process | 10 | 3 | 0.64 | 0.02197 |
| GO:0043288 | apocarotenoid metabolic process | 10 | 3 | 0.64 | 0.02197 |
| GO:0009687 | abscisic acid metabolic process | 10 | 3 | 0.64 | 0.02197 |
| GO:0046474 | glycerophospholipid biosynthetic process | 26 | 5 | 1.67 | 0.02198 |
| GO:0023056 | positive regulation of signaling | 26 | 5 | 1.67 | 0.02198 |
| GO:0010647 | positive regulation of cell communicatio... | 26 | 5 | 1.67 | 0.02198 |
| GO:0009967 | positive regulation of signal transductiion | 26 | 5 | 1.67 | 0.02198 |
| GO:0009409 | response to cold | 112 | 13 | 7.18 | 0.02407 |
| GO:0032509 | endosome transport via multivesicular | 18 | 4 | 1.15 | 0.02422 |

**Functions enriched among genes that are down-regulated in *A. nemorensis* but up regulated in *A. sagittata* during wilting.**

| GO.ID | Term | Annotated | Significant | Expected | KS |
|---|---|---|---|---|---|

| | | | | | |
|---|---|---|---|---|---|
| GO:0009751 | response to salicylic acid | 25 | 11 | 2.62 | 1.6e-05 |
| GO:0002237 | response to molecule of bacterial origin | 10 | 6 | 1.05 | 0.00018 |
| GO:0071456 | cellular response to hypoxia | 22 | 9 | 2.31 | 0.0002 |
| GO:0009755 | hormone-mediated signaling pathway | 119 | 29 | 12.49 | 0.0005 |
| GO:0009611 | response to wounding | 56 | 14 | 5.88 | 0.0014 |
| GO:0009735 | response to cytokinin | 18 | 7 | 1.89 | 0.00149 |
| GO:0010039 | response to iron ion | 10 | 5 | 1.05 | 0.00199 |
| GO:0071407 | cellular response to organic cyclic compound | 30 | 9 | 3.15 | 0.00265 |
| GO:0009813 | flavonoid biosynthetic process | 15 | 6 | 1.57 | 0.00279 |
| GO:0009414 | response to water deprivation | 93 | 19 | 9.76 | 0.00284 |
| GO:0009738 | abscisic acid-activated signaling pathwa... | 36 | 10 | 3.78 | 0.00293 |
| GO:0009555 | pollen development | 42 | 11 | 4.41 | 0.00304 |
| GO:0016567 | protein ubiquitination | 80 | 20 | 8.4 | 0.00316 |
| GO:0019748 | secondary metabolic process | 43 | 11 | 4.51 | 0.00372 |
| GO:0006511 | ubiquitin-dependent protein catabolic pr... | 56 | 13 | 5.88 | 0.00417 |
| GO:0045088 | regulation of innate immune response | 38 | 10 | 3.99 | 0.0045 |
| GO:0000209 | protein polyubiquitination | 12 | 5 | 1.26 | 0.00523 |
| GO:0009251 | glucan catabolic process | 12 | 5 | 1.26 | 0.00523 |
| GO:0009626 | plant-type hypersensitive response | 17 | 6 | 1.78 | 0.00576 |
| GO:0042742 | defense response to bacterium | 78 | 16 | 8.19 | 0.00577 |
| GO:0036211 | protein modification process | 263 | 49 | 27.6 | 0.00706 |
| GO:0031349 | positive regulation of defense response | 23 | 7 | 2.41 | 0.00725 |
| GO:0043067 | regulation of programmed cell death | 18 | 6 | 1.89 | 0.00789 |
| GO:0010468 | regulation of gene expression | 290 | 43 | 30.44 | 0.00907 |
| GO:0000041 | transition metal ion transport | 24 | 7 | 2.52 | 0.00934 |
| GO:0009615 | response to virus | 19 | 6 | 1.99 | 0.01054 |
| GO:0043207 | response to external biotic stimulus | 222 | 45 | 23.3 | 0.01123 |

| | | | | | |
|---|---|---|---|---|---|
| GO:0051707 | response to other organism | 222 | 45 | 23.3 | 0.01123 |
| GO:0009607 | response to biotic stimulus | 223 | 45 | 23.41 | 0.01233 |
| GO:0006355 | regulation of DNA-templated transcription | 206 | 32 | 21.62 | 0.01246 |

**Functions enriched among down-regulated genes that are down regulated to a lower level in *A. sagittata* as compared to *A. nemorensis* in wilting.**

| GO.ID | Term | Annotated | Significant | Expected | KS |
|---|---|---|---|---|---|
| GO:0006412 | translation | 164 | 62 | 19.65 | 3.0e-14 |
| GO:0045037 | protein import into chloroplast stroma | 14 | 9 | 1.68 | 5.3e-06 |
| GO:0009658 | chloroplast organization | 81 | 24 | 9.71 | 1.2e-05 |
| GO:0009793 | embryo development ending in seed dormancy | 104 | 25 | 12.46 | 0.00036 |
| GO:0006783 | heme biosynthetic process | 10 | 6 | 1.2 | 0.00039 |
| GO:0002181 | cytoplasmic translation | 18 | 8 | 2.16 | 0.00058 |
| GO:0006364 | rRNA processing | 31 | 13 | 3.72 | 0.00083 |
| GO:0072596 | establishment of protein localization | 26 | 15 | 3.12 | 0.00122 |
| GO:0009657 | plastid organization | 112 | 34 | 13.42 | 0.0016 |
| GO:0006418 | tRNA aminoacylation for protein translation | 21 | 8 | 2.52 | 0.00194 |
| GO:0015995 | chlorophyll biosynthetic process | 37 | 11 | 4.43 | 0.00294 |
| GO:0071806 | protein transmembrane transport | 24 | 14 | 2.88 | 0.00325 |
| GO:0010410 | hemicellulose metabolic process | 14 | 6 | 1.68 | 0.00366 |
| GO:0140053 | mitochondrial gene expression | 11 | 5 | 1.32 | 0.00599 |
| GO:0090150 | establishment of protein localization | 20 | 7 | 2.4 | 0.00643 |
| GO:0043648 | dicarboxylic acid metabolic process | 36 | 10 | 4.31 | 0.00768 |
| GO:0016116 | carotenoid metabolic process | 21 | 7 | 2.52 | 0.00867 |
| GO:0051085 | chaperone cofactor-dependent protein | 12 | 5 | 1.44 | 0.00928 |
| GO:1901259 | chloroplast rRNA processing | 12 | 5 | 1.44 | 0.00928 |
| GO:0042026 | protein refolding | 12 | 5 | 1.44 | 0.00928 |
| GO:0042254 | ribosome biogenesis | 54 | 20 | 6.47 | 0.0126 |
| GO:0010027 | thylakoid membrane organization | 28 | 8 | 3.36 | 0.01395 |

| | | | | | |
|---|---|---|---|---|---|
| GO:0009668 | plastid membrane organization | 28 | 8 | 3.36 | 0.01395 |
| GO:0019684 | photosynthesis, light reaction | 76 | 16 | 9.11 | 0.0156 |
| GO:0022613 | ribonucleoprotein complex biogenesis | 55 | 20 | 6.59 | 0.01607 |
| GO:0009073 | aromatic amino acid family biosynthetic process | 25 | 7 | 3 | 0.02352 |
| GO:2000070 | regulation of response to water deprivation | 10 | 4 | 1.2 | 0.02359 |

**Functions enriched among up-regulated genes that were less up-regulated in *A. sagittata* as compared to *A. nemorensis* during wilting.**

| GO.ID | Term | Annotated | Significant | Expected | KS |
|---|---|---|---|---|---|
| GO:0071489 | cellular response to red or far red light | 15 | 5 | 0.46 | 0.00087 |
| GO:0042026 | protein refolding | 18 | 4 | 0.55 | 0.00179 |
| GO:1901615 | organic hydroxy compound metabolic process | 67 | 7 | 2.05 | 0.00382 |
| GO:0044282 | small molecule catabolic process | 67 | 7 | 2.05 | 0.00382 |
| GO:0061077 | chaperone-mediated protein folding | 22 | 4 | 0.67 | 0.0039 |
| GO:0019748 | secondary metabolic process | 37 | 7 | 1.13 | 0.00414 |
| GO:0034605 | cellular response to heat | 24 | 4 | 0.73 | 0.0054 |
| GO:0010017 | red or far-red light signaling pathway | 13 | 3 | 0.4 | 0.0063 |
| GO:0009408 | response to heat | 82 | 10 | 2.5 | 0.00643 |
| GO:0030003 | intracellular monoatomic cation homeostasis | 41 | 5 | 1.25 | 0.00746 |
| GO:0009809 | lignin biosynthetic process | 14 | 3 | 0.43 | 0.00785 |
| GO:0006720 | isoprenoid metabolic process | 42 | 5 | 1.28 | 0.00827 |
| GO:0120255 | olefinic compound biosynthetic process | 15 | 3 | 0.46 | 0.00959 |
| GO:0019752 | carboxylic acid metabolic process | 210 | 13 | 6.41 | 0.01011 |
| GO:0051084 | 'de novo' post-translational protein fol... | 17 | 3 | 0.52 | 0.01372 |
| GO:0051085 | chaperone cofactor-dependent protein | 17 | 3 | 0.52 | 0.01372 |
| GO:0006721 | terpenoid metabolic process | 32 | 4 | 0.98 | 0.01519 |

| | | | | | |
|---|---|---|---|---|---|
| GO:0006979 | response to oxidative stress | 87 | 7 | 2.66 | 0.01569 |
| GO:0006458 | 'de novo' protein folding | 18 | 3 | 0.55 | 0.01611 |
| GO:0009812 | flavonoid metabolic process | 18 | 3 | 0.55 | 0.01611 |
| GO:0009699 | phenylpropanoid biosynthetic process | 21 | 5 | 0.64 | 0.01656 |
| GO:0006082 | organic acid metabolic process | 224 | 13 | 6.84 | 0.01682 |
| GO:0043436 | oxoacid metabolic process | 224 | 13 | 6.84 | 0.01682 |
| GO:0044248 | cellular catabolic process | 132 | 9 | 4.03 | 0.01783 |
| GO:0016054 | organic acid catabolic process | 51 | 5 | 1.56 | 0.01842 |
| GO:0046395 | carboxylic acid catabolic process | 51 | 5 | 1.56 | 0.01842 |
| GO:0006520 | amino acid metabolic process | 90 | 7 | 2.75 | 0.01864 |
| GO:0044283 | small molecule biosynthetic process | 135 | 9 | 4.12 | 0.02038 |
| GO:0098660 | inorganic ion transmembrane transport | 53 | 5 | 1.62 | 0.02146 |
| GO:0044550 | secondary metabolite biosynthetic process | 22 | 5 | 0.67 | 0.02166 |

**Functions enriched among genes that responded in opposite ways to wilting, up regulated in *A. nemorensis* and down regulated in *A. sagittata***

| GO.ID | Term | Annotated | Significant | Expected | KS |
|---|---|---|---|---|---|
| GO:0009658 | chloroplast organization | 53 | 20 | 5.85 | 2.7e-07 |
| GO:0006413 | translational initiation | 35 | 13 | 3.87 | 4.4e-05 |
| GO:0006412 | translation | 110 | 36 | 12.15 | 5.2e-05 |
| GO:0010027 | thylakoid membrane organization | 10 | 6 | 1.1 | 0.00025 |
| GO:0045037 | protein import into chloroplast stroma | 10 | 6 | 1.1 | 0.00025 |
| GO:0046364 | monosaccharide biosynthetic process | 11 | 6 | 1.22 | 0.00049 |
| GO:0042274 | ribosomal small subunit biogenesis | 21 | 8 | 2.32 | 0.00113 |
| GO:0006099 | tricarboxylic acid cycle | 13 | 6 | 1.44 | 0.00151 |
| GO:0009793 | embryo development ending in seed dormancy | 148 | 28 | 16.35 | 0.00249 |
| GO:0006753 | nucleoside phosphate metabolic process | 93 | 23 | 10.27 | 0.00261 |
| GO:0010109 | regulation of photosynthesis | 16 | 6 | 1.77 | 0.0053 |

| | | | | | |
|---|---|---|---|---|---|
| GO:0006364 | rRNA processing | 61 | 14 | 6.74 | 0.00539 |
| GO:0022900 | electron transport chain | 21 | 7 | 2.32 | 0.00554 |
| GO:0006414 | translational elongation | 12 | 5 | 1.33 | 0.00654 |
| GO:0051156 | glucose 6-phosphate metabolic process | 12 | 5 | 1.33 | 0.00654 |
| GO:0006096 | glycolytic process | 17 | 6 | 1.88 | 0.00744 |
| GO:1901137 | carbohydrate derivative biosynthetic process | 70 | 15 | 7.73 | 0.00791 |
| GO:0030244 | cellulose biosynthetic process | 23 | 7 | 2.54 | 0.00961 |
| GO:0072596 | establishment of protein localization | 12 | 8 | 1.33 | 0.0118 |
| GO:0044283 | small molecule biosynthetic process | 135 | 28 | 14.91 | 0.01273 |
| GO:0006767 | water-soluble vitamin metabolic process | 14 | 5 | 1.55 | 0.01373 |
| GO:0006790 | sulfur compound metabolic process | 68 | 14 | 7.51 | 0.01442 |
| GO:0002181 | cytoplasmic translation | 25 | 7 | 2.76 | 0.01553 |
| GO:0019693 | ribose phosphate metabolic process | 37 | 12 | 4.09 | 0.0165 |
| GO:0006006 | glucose metabolic process | 10 | 4 | 1.1 | 0.01786 |
| GO:0072527 | pyrimidine-containing compound metabolic process | 10 | 4 | 1.1 | 0.01786 |
| GO:0065003 | protein-containing complex assembly | 119 | 21 | 13.15 | 0.01835 |
| GO:0046394 | carboxylic acid biosynthetic process | 98 | 18 | 10.83 | 0.01913 |
| GO:0016053 | organic acid biosynthetic process | 98 | 18 | 10.83 | 0.01913 |
| GO:0009698 | phenylpropanoid metabolic process | 26 | 7 | 2.87 | 0.0193 |

**Functions enriched among down-regulated genes that were less down-regulated in *A. sagittata* as compared to *A. nemorensis* during wilting.**

| GO.ID | Term | Annotated | Significant | Expected | KS |
|---|---|---|---|---|---|
| GO:0006355 | regulation of DNA-templated transcription | 206 | 23 | 11.32 | 0.00062 |
| GO:0045087 | innate immune response | 59 | 10 | 3.24 | 0.00113 |
| GO:0016310 | phosphorylation | 120 | 15 | 6.6 | 0.00192 |
| GO:0009624 | response to nematode | 20 | 5 | 1.1 | 0.00371 |
| GO:0019761 | glucosinolate biosynthetic process | 20 | 5 | 1.1 | 0.00371 |

| | | | | | |
|---|---|---|---|---|---|
| GO:0000103 | sulfate assimilation | 13 | 4 | 0.71 | 0.00424 |
| GO:0050776 | regulation of immune response | 41 | 7 | 2.25 | 0.00608 |
| GO:0031348 | negative regulation of defense response | 23 | 5 | 1.26 | 0.00704 |

**Table-S5:** Genes showing a different change in expression level after recovery in *A. sagittata* compared to *A. nemorensis* at FDR<0.05 are enriched in several biological functional GO categories. Given are GO id number, term description, number of annotated genes in this category, number of genes responding more in *A. sagittata* that belong to this category, expected number of genes, *p*-value of the enrichment. Enrichment were computed separately depending on the pattern of differential expression.

**Functions enriched among genes that respond similarly in both *A. sagittata* and *A. nemorensis* after recovery**

| GO.ID | Term | Annotated | Significant | Expected | KS |
|---|---|---|---|---|---|
| GO:0002237 | response to molecule of bacterial origin | 9 | 5 | 0.74 | 0.00034 |
| GO:0009813 | flavonoid biosynthetic process | 18 | 6 | 1.48 | 0.00228 |
| GO:0080134 | regulation of response to stress | 92 | 16 | 7.55 | 0.00272 |
| GO:0046283 | anthocyanin-containing compound metaboli... | 14 | 5 | 1.15 | 0.00385 |
| GO:0010182 | sugar mediated signaling pathway | 9 | 4 | 0.74 | 0.004 |
| GO:0051716 | cellular response to stimulus | 528 | 61 | 43.32 | 0.00588 |
| GO:0009651 | response to salt stress | 126 | 19 | 10.34 | 0.00597 |
| GO:0010311 | lateral root formation | 16 | 5 | 1.31 | 0.00733 |
| GO:0009751 | response to salicylic acid | 30 | 7 | 2.46 | 0.00903 |
| GO:0009819 | drought recovery | 6 | 3 | 0.49 | 0.00905 |

| | | | | | |
|---|---|---|---|---|---|
| GO:0009415 | response to water | 111 | 19 | 9.11 | 0.00925 |
| GO:0009611 | response to wounding | 45 | 9 | 3.69 | 0.00929 |
| GO:0031669 | cellular response to nutrient levels | 47 | 9 | 3.86 | 0.01237 |
| GO:1902680 | positive regulation of RNA biosynthetic ... | 72 | 12 | 5.91 | 0.01269 |
| GO:0045893 | positive regulation of DNA-templated tra... | 72 | 12 | 5.91 | 0.01269 |

**Functions enriched among up-regulated genes that were more up-regulated in in *A. sagittata* than in *A. nemorensis* after recovery**

| GO.ID | Term | Annotated | Significant | Expected | KS |
|---|---|---|---|---|---|
| GO:0005982 | starch metabolic process | 25 | 7 | 1.11 | 7.00E-05 |
| GO:1901700 | response to oxygen-containing compound | 349 | 29 | 15.48 | 0.00042 |
| GO:0033993 | response to lipid | 197 | 18 | 8.74 | 0.00216 |
| GO:0043603 | amide metabolic process | 44 | 7 | 1.95 | 0.00277 |
| GO:0010876 | lipid localization | 16 | 4 | 0.71 | 0.00443 |
| GO:0015711 | organic anion transport | 16 | 4 | 0.71 | 0.00443 |
| GO:0009408 | response to heat | 74 | 9 | 3.28 | 0.00477 |
| GO:0009615 | response to virus | 26 | 5 | 1.15 | 0.00491 |
| GO:0071456 | cellular response to hypoxia | 28 | 5 | 1.24 | 0.00684 |
| GO:0048868 | pollen tube development | 20 | 4 | 0.89 | 0.01028 |
| GO:0009737 | response to abscisic acid | 129 | 12 | 5.72 | 0.01058 |
| GO:0097305 | response to alcohol | 132 | 12 | 5.86 | 0.0126 |

**Functions enriched among genes that were up regulated in *A. sagittata* but down regulated in *A. nemorensis* after recovery**

| GO.ID | Term | Annotated | Significant | Expected | KS |
|---|---|---|---|---|---|

| | | | | | |
|---|---|---|---|---|---|
| GO:0009414 | response to water deprivation | 103 | 27 | 13.49 | 0.0002 |
| GO:0005982 | starch metabolic process | 27 | 10 | 3.54 | 0.0014 |
| GO:0010119 | regulation of stomatal movement | 36 | 11 | 4.71 | 0.0047 |
| GO:0051239 | regulation of multicellular organismal process | 92 | 24 | 12.05 | 0.0048 |
| GO:0009749 | response to glucose | 18 | 7 | 2.36 | 0.0054 |
| GO:0007389 | pattern specification process | 42 | 12 | 5.5 | 0.0059 |
| GO:2000026 | regulation of multicellular organism | 80 | 19 | 10.48 | 0.0059 |
| GO:0006970 | response to osmotic stress | 133 | 28 | 17.42 | 0.0059 |
| GO:0051240 | positive regulation of multicellular organization | 28 | 9 | 3.67 | 0.0072 |
| GO:0050832 | defense response to fungus | 43 | 12 | 5.63 | 0.0072 |
| GO:0016567 | protein ubiquitination | 93 | 21 | 12.18 | 0.0073 |
| GO:0051603 | proteolysis involved in protein catabolic process | 130 | 27 | 17.03 | 0.0083 |
| GO:0009789 | positive regulation of abscisic acid-activation | 15 | 6 | 1.96 | 0.0086 |
| GO:0000272 | polysaccharide catabolic process | 39 | 11 | 5.11 | 0.0091 |
| GO:0097305 | response to alcohol | 123 | 29 | 16.11 | 0.0093 |
| GO:0006914 | autophagy | 34 | 10 | 4.45 | 0.0093 |
| GO:0048580 | regulation of post-embryonic development | 79 | 18 | 10.35 | 0.0114 |

| | | | | | |
|---|---|---|---|---|---|
| GO:0090693 | plant organ senescence | 25 | 8 | 3.27 | 0.0114 |
| GO:0048582 | positive regulation of post-embryonic | 25 | 8 | 3.27 | 0.0114 |
| GO:0010150 | leaf senescence | 25 | 8 | 3.27 | 0.0114 |
| GO:0009737 | response to abscisic acid | 119 | 28 | 15.58 | 0.0117 |
| GO:0042594 | response to starvation | 41 | 11 | 5.37 | 0.0135 |
| GO:0016236 | macroautophagy | 21 | 7 | 2.75 | 0.014 |
| GO:0050896 | response to stimulus | 1064 | 175 | 139.35 | 0.0182 |
| GO:0009651 | response to salt stress | 108 | 22 | 14.14 | 0.0204 |
| GO:0071215 | cellular response to abscisic acid stimulus | 48 | 15 | 6.29 | 0.0207 |
| GO:0097306 | cellular response to alcohol | 48 | 15 | 6.29 | 0.0207 |
| GO:0009617 | response to bacterium | 115 | 23 | 15.06 | 0.0221 |
| GO:0051606 | detection of stimulus | 18 | 6 | 2.36 | 0.0226 |

**Functions enriched among down-regulated genes that were down-regulated at an even lower level in *A. sagittata* as compared to *A. nemorensis* after recovery**

| GO.ID | Term | Annotated | Significant | Expected | KS |
|---|---|---|---|---|---|
| GO:0006006 | glucose metabolic process | 23 | 10 | 1.16 | 5.2e-08 |
| GO:0015979 | photosynthesis | 93 | 19 | 4.71 | 8.4e-08 |
| GO:0019319 | hexose biosynthetic process | 15 | 6 | 0.76 | 5.2e-05 |
| GO:0015995 | chlorophyll biosynthetic process | 23 | 7 | 1.16 | 9.1e-05 |
| GO:0006417 | regulation of translation | 17 | 6 | 0.86 | 0.00012 |
| GO:0006096 | glycolytic process | 18 | 6 | 0.91 | 0.00017 |

**Functions enriched among up-regulated genes that were less up-regulated in *A. sagittata* than in *A. nemorensis* in recovery**

| GO.ID | Term | Annotated | Significant | Expected | KS |
|---|---|---|---|---|---|
| GO:0042221 | response to chemical | 536 | 12 | 6.19 | 0.012 |

| | | | | | |
|---|---|---|---|---|---|
| GO:0006790 | sulfur compound metabolic process | 81 | 4 | 0.93 | 0.013 |
| GO:0046686 | response to cadmium ion | 16 | 2 | 0.18 | 0.014 |
| GO:0050896 | response to stimulus | 1076 | 19 | 12.42 | 0.014 |
| GO:0042254 | ribosome biogenesis | 128 | 5 | 1.48 | 0.014 |
| GO:0016070 | RNA metabolic process | 618 | 13 | 7.13 | 0.015 |
| GO:0010038 | response to metal ion | 46 | 3 | 0.53 | 0.015 |
| GO:0010109 | regulation of photosynthesis | 18 | 2 | 0.21 | 0.018 |
| GO:0006396 | RNA processing | 247 | 7 | 2.85 | 0.02 |
| GO:0006109 | regulation of carbohydrate metabolic pro... | 22 | 2 | 0.25 | 0.026 |
| GO:0022613 | ribonucleoprotein complex biogenesis | 153 | 5 | 1.77 | 0.029 |
| GO:0090304 | nucleic acid metabolic process | 683 | 13 | 7.88 | 0.033 |
| GO:0062012 | regulation of small molecule metabolic p... | 26 | 2 | 0.3 | 0.035 |
| GO:0010467 | gene expression | 922 | 16 | 10.64 | 0.036 |
| GO:0044249 | cellular biosynthetic process | 1174 | 19 | 13.55 | 0.038 |
| GO:0032774 | RNA biosynthetic process | 551 | 11 | 6.36 | 0.038 |
| GO:0141187 | nucleic acid biosynthetic process | 554 | 11 | 6.39 | 0.04 |
| GO:0009751 | response to salicylic acid | 30 | 2 | 0.35 | 0.046 |
| GO:0042273 | ribosomal large subunit biogenesis | 30 | 2 | 0.35 | 0.046 |
| GO:0009739 | response to gibberellin | 31 | 2 | 0.36 | 0.049 |

| | | | | | |
|---|---|---|---|---|---|
| GO:0006418 | tRNA aminoacylation for protein translat... | 31 | 2 | 0.36 | 0.049 |
| GO:0043039 | tRNA aminoacylation | 32 | 2 | 0.37 | 0.052 |
| GO:0043038 | amino acid activation | 33 | 2 | 0.38 | 0.055 |
| GO:0010150 | leaf senescence | 33 | 2 | 0.38 | 0.055 |
| GO:0090693 | plant organ senescence | 34 | 2 | 0.39 | 0.058 |
| GO:0009059 | macromolecule biosynthetic process | 982 | 16 | 11.33 | 0.063 |
| GO:0034654 | nucleobase-containing compound biosynthe... | 597 | 11 | 6.89 | 0.064 |
| GO:0019684 | photosynthesis, light reaction | 38 | 2 | 0.44 | 0.07 |
| GO:0002181 | cytoplasmic translation | 39 | 2 | 0.45 | 0.073 |
| GO:0006364 | rRNA processing | 87 | 3 | 1 | 0.077 |

**Functions enriched among genes that were up regulated in *A. nemorensis* but down regulated in *A. sagittata* after recovery**

| GO.ID | Term | Annotated | Significant | Expected | KS |
|---|---|---|---|---|---|
| GO:0006412 | translation | 230 | 138 | 39.31 | < 1e-30 |
| GO:0002181 | cytoplasmic translation | 39 | 24 | 6.67 | 4.6e-10 |
| GO:0009658 | chloroplast organization | 90 | 33 | 15.38 | 4.7e-06 |
| GO:0042255 | ribosome assembly | 18 | 11 | 3.08 | 3.3e-05 |
| GO:0006414 | translational elongation | 18 | 11 | 3.08 | 3.3e-05 |
| GO:0010027 | thylakoid membrane organization | 23 | 12 | 3.93 | 0.00012 |
| GO:0046034 | ATP metabolic process | 17 | 10 | 2.91 | 0.00012 |
| GO:0042274 | ribosomal small subunit biogenesis | 34 | 15 | 5.81 | 0.00019 |

| | | | | | |
|---|---|---|---|---|---|
| GO:0045037 | protein import into chloroplast stroma | 18 | 10 | 3.08 | 0.00023 |
| GO:0015995 | chlorophyll biosynthetic process | 26 | 12 | 4.44 | 0.00052 |
| GO:0019318 | hexose metabolic process | 20 | 10 | 3.42 | 0.0007 |
| GO:0046434 | organophosphate catabolic process | 18 | 9 | 3.08 | 0.00131 |
| GO:0019684 | photosynthesis, light reaction | 38 | 14 | 6.5 | 0.00267 |
| GO:0006090 | pyruvate metabolic process | 20 | 9 | 3.42 | 0.00328 |
| GO:0065003 | protein-containing complex assembly | 124 | 33 | 21.2 | 0.00426 |
| GO:0015979 | photosynthesis | 59 | 23 | 10.09 | 0.00436 |
| GO:0072596 | establishment of protein localization to... | 29 | 16 | 4.96 | 0.00478 |
| GO:0051085 | chaperone cofactor-dependent protein ref... | 15 | 7 | 2.56 | 0.00745 |
| GO:0072526 | pyridine-containing compound catabolic p... | 15 | 7 | 2.56 | 0.00745 |
| GO:0022900 | electron transport chain | 26 | 10 | 4.44 | 0.0076 |
| GO:0042273 | ribosomal large subunit biogenesis | 30 | 11 | 5.13 | 0.00785 |
| GO:0018193 | peptidyl-amino acid modification | 19 | 8 | 3.25 | 0.00898 |
| GO:0006413 | translational initiation | 31 | 11 | 5.3 | 0.01035 |
| GO:0009078 | pyruvate family amino acid metabolic pro... | 16 | 7 | 2.73 | 0.01134 |
| GO:0009260 | ribonucleotide biosynthetic process | 20 | 8 | 3.42 | 0.01277 |

| | | | | | |
|---|---|---|---|---|---|
| GO:0009657 | plastid organization | 118 | 45 | 20.17 | 0.01284 |
| GO:0042254 | ribosome biogenesis | 128 | 44 | 21.88 | 0.01286 |

**Functions enriched among down-regulated genes that were less down-regulated in *A. sagittata* as compared to *A. nemorensis* after recovery**

| GO.ID | Term | Annotated | Significant | Expected | KS |
|---|---|---|---|---|---|
| GO:0036294 | cellular response to decreased oxygen le... | 28 | 2 | 0.2 | 0.017 |
| GO:0071453 | cellular response to oxygen levels | 28 | 2 | 0.2 | 0.017 |
| GO:0071456 | cellular response to hypoxia | 28 | 2 | 0.2 | 0.017 |
| GO:0015711 | organic anion transport | 32 | 2 | 0.23 | 0.021 |
| GO:0001666 | response to hypoxia | 36 | 2 | 0.26 | 0.027 |
| GO:0036293 | response to decreased oxygen levels | 36 | 2 | 0.26 | 0.027 |
| GO:0070482 | response to oxygen levels | 36 | 2 | 0.26 | 0.027 |
| GO:0006873 | intracellular monoatomic ion homeostasis | 37 | 2 | 0.27 | 0.028 |

**Table-S6:** The performance of models was evaluated by loss, accuracy, auROC and auPR deepCRE-like (Peleke et al., 2024) models trained on RNAseq data of *Arabis nemorensis* and *A. sagittata* treatments control, wilting and recovery.

| Species | Treatment | Training-Strategy | Chromosome | Loss | Accuracy | auROC | auPR |
|---|---|---|---|---|---|---|---|
| *Arabis nemorensis* | control | SSC | 1 | 0.653 | 0.608 | 0.650 | 0.610 |
| *Arabis nemorensis* | control | SSC | 2 | 0.661 | 0.600 | 0.618 | 0.579 |

| | | | | | | | |
|---|---|---|---|---|---|---|---|
| *Arabis nemorensis* | control | SSC | 3 | 0.652 | 0.614 | 0.636 | 0.572 |
| *Arabis nemorensis* | control | SSC | 4 | 0.651 | 0.598 | 0.653 | 0.617 |
| *Arabis nemorensis* | control | SSC | 5 | 0.652 | 0.591 | 0.628 | 0.592 |
| *Arabis nemorensis* | control | SSC | 6 | 0.666 | 0.592 | 0.613 | 0.591 |
| *Arabis nemorensis* | control | SSC | 7 | 0.660 | 0.591 | 0.612 | 0.584 |
| *Arabis nemorensis* | control | SSC | 8 | 0.661 | 0.582 | 0.627 | 0.597 |
| *Arabis nemorensis* | control | SSR | 1 | 0.529 | 0.741 | 0.815 | 0.805 |
| *Arabis nemorensis* | control | SSR | 2 | 0.513 | 0.768 | 0.831 | 0.812 |
| *Arabis nemorensis* | control | SSR | 3 | 0.514 | 0.757 | 0.829 | 0.805 |
| *Arabis nemorensis* | control | SSR | 4 | 0.512 | 0.755 | 0.832 | 0.809 |
| *Arabis nemorensis* | control | SSR | 5 | 0.479 | 0.769 | 0.855 | 0.847 |
| *Arabis nemorensis* | control | SSR | 6 | 0.521 | 0.757 | 0.827 | 0.814 |
| *Arabis nemorensis* | control | SSR | 7 | 0.513 | 0.763 | 0.829 | 0.824 |
| *Arabis nemorensis* | control | SSR | 8 | 0.522 | 0.755 | 0.822 | 0.802 |
| *Arabis nemorensis* | recovery | SSC | 1 | 0.657 | 0.583 | 0.599 | 0.556 |

| | | | | | | | |
|---|---|---|---|---|---|---|---|
| *Arabis nemorensis* | recovery | SSC | 2 | 0.663 | 0.583 | 0.623 | 0.600 |
| *Arabis nemorensis* | recovery | SSC | 3 | 0.646 | 0.632 | 0.667 | 0.593 |
| *Arabis nemorensis* | recovery | SSC | 4 | 0.639 | 0.624 | 0.681 | 0.639 |
| *Arabis nemorensis* | recovery | SSC | 5 | 0.646 | 0.615 | 0.671 | 0.642 |
| *Arabis nemorensis* | recovery | SSC | 6 | 0.660 | 0.592 | 0.638 | 0.592 |
| *Arabis nemorensis* | recovery | SSC | 7 | 0.662 | 0.586 | 0.650 | 0.618 |
| *Arabis nemorensis* | recovery | SSC | 8 | 0.645 | 0.613 | 0.675 | 0.637 |
| *Arabis nemorensis* | recovery | SSR | 1 | 0.536 | 0.742 | 0.811 | 0.797 |
| *Arabis nemorensis* | recovery | SSR | 2 | 0.509 | 0.764 | 0.831 | 0.801 |
| *Arabis nemorensis* | recovery | SSR | 3 | 0.505 | 0.767 | 0.842 | 0.829 |
| *Arabis nemorensis* | recovery | SSR | 4 | 0.503 | 0.752 | 0.839 | 0.819 |
| *Arabis nemorensis* | recovery | SSR | 5 | 0.493 | 0.775 | 0.846 | 0.828 |
| *Arabis nemorensis* | recovery | SSR | 6 | 0.512 | 0.758 | 0.829 | 0.823 |
| *Arabis nemorensis* | recovery | SSR | 7 | 0.505 | 0.764 | 0.837 | 0.836 |
| *Arabis nemorensis* | recovery | SSR | 8 | 0.500 | 0.761 | 0.840 | 0.821 |

| | | | | | | | |
|---|---|---|---|---|---|---|---|
| *Arabis nemorensis* | wilting | SSC | 1 | 0.646 | 0.610 | 0.648 | 0.603 |
| *Arabis nemorensis* | wilting | SSC | 2 | 0.649 | 0.620 | 0.666 | 0.641 |
| *Arabis nemorensis* | wilting | SSC | 3 | 0.646 | 0.621 | 0.661 | 0.592 |
| *Arabis nemorensis* | wilting | SSC | 4 | 0.642 | 0.624 | 0.692 | 0.655 |
| *Arabis nemorensis* | wilting | SSC | 5 | 0.651 | 0.599 | 0.646 | 0.596 |
| *Arabis nemorensis* | wilting | SSC | 6 | 0.656 | 0.617 | 0.655 | 0.622 |
| *Arabis nemorensis* | wilting | SSC | 7 | 0.649 | 0.622 | 0.661 | 0.620 |
| *Arabis nemorensis* | wilting | SSC | 8 | 0.646 | 0.604 | 0.677 | 0.636 |
| *Arabis nemorensis* | wilting | SSR | 1 | 0.522 | 0.755 | 0.825 | 0.807 |
| *Arabis nemorensis* | wilting | SSR | 2 | 0.512 | 0.753 | 0.827 | 0.804 |
| *Arabis nemorensis* | wilting | SSR | 3 | 0.499 | 0.769 | 0.842 | 0.829 |
| *Arabis nemorensis* | wilting | SSR | 4 | 0.493 | 0.765 | 0.844 | 0.829 |
| *Arabis nemorensis* | wilting | SSR | 5 | 0.497 | 0.769 | 0.844 | 0.824 |
| *Arabis nemorensis* | wilting | SSR | 6 | 0.481 | 0.778 | 0.853 | 0.836 |
| *Arabis nemorensis* | wilting | SSR | 7 | 0.491 | 0.773 | 0.845 | 0.845 |

| | | | | | | | |
|---|---|---|---|---|---|---|---|
| *Arabis nemorensis* | wilting | SSR | 8 | 0.525 | 0.746 | 0.822 | 0.793 |
| *Arabis sagittata* | control | SSC | chr1 | 0.657 | 0.586 | 0.639 | 0.611 |
| *Arabis sagittata* | control | SSC | chr2 | 0.644 | 0.624 | 0.674 | 0.635 |
| *Arabis sagittata* | control | SSC | chr3 | 0.657 | 0.596 | 0.618 | 0.579 |
| *Arabis sagittata* | control | SSC | chr4 | 0.642 | 0.607 | 0.683 | 0.647 |
| *Arabis sagittata* | control | SSC | chr5 | 0.657 | 0.582 | 0.639 | 0.605 |
| *Arabis sagittata* | control | SSC | chr6 | 0.654 | 0.601 | 0.640 | 0.609 |
| *Arabis sagittata* | control | SSC | chr7 | 0.663 | 0.579 | 0.642 | 0.607 |
| *Arabis sagittata* | control | SSC | chr8 | 0.656 | 0.596 | 0.640 | 0.615 |
| *Arabis sagittata* | control | SSR | chr1 | 0.545 | 0.724 | 0.799 | 0.777 |
| *Arabis sagittata* | control | SSR | chr2 | 0.499 | 0.775 | 0.838 | 0.821 |
| *Arabis sagittata* | control | SSR | chr3 | 0.528 | 0.750 | 0.817 | 0.796 |
| *Arabis sagittata* | control | SSR | chr4 | 0.503 | 0.757 | 0.837 | 0.822 |
| *Arabis sagittata* | control | SSR | chr5 | 0.485 | 0.780 | 0.849 | 0.832 |
| *Arabis sagittata* | control | SSR | chr6 | 0.510 | 0.747 | 0.830 | 0.807 |

| | | | | | | | |
|---|---|---|---|---|---|---|---|
| *Arabis sagittata* | control | SSR | chr7 | 0.520 | 0.753 | 0.826 | 0.822 |
| *Arabis sagittata* | control | SSR | chr8 | 0.531 | 0.732 | 0.811 | 0.785 |
| *Arabis sagittata* | recovery | SSC | chr1 | 0.657 | 0.597 | 0.639 | 0.604 |
| *Arabis sagittata* | recovery | SSC | chr2 | 0.660 | 0.598 | 0.627 | 0.586 |
| *Arabis sagittata* | recovery | SSC | chr3 | 0.645 | 0.612 | 0.659 | 0.624 |
| *Arabis sagittata* | recovery | SSC | chr4 | 0.638 | 0.629 | 0.678 | 0.634 |
| *Arabis sagittata* | recovery | SSC | chr5 | 0.654 | 0.608 | 0.627 | 0.579 |
| *Arabis sagittata* | recovery | SSC | chr6 | 0.656 | 0.603 | 0.650 | 0.621 |
| *Arabis sagittata* | recovery | SSC | chr7 | 0.665 | 0.591 | 0.638 | 0.599 |
| *Arabis sagittata* | recovery | SSC | chr8 | 0.661 | 0.589 | 0.620 | 0.587 |
| *Arabis sagittata* | recovery | SSR | chr1 | 0.520 | 0.755 | 0.825 | 0.807 |
| *Arabis sagittata* | recovery | SSR | chr2 | 0.526 | 0.750 | 0.819 | 0.806 |
| *Arabis sagittata* | recovery | SSR | chr3 | 0.501 | 0.771 | 0.840 | 0.817 |
| *Arabis sagittata* | recovery | SSR | chr4 | 0.516 | 0.748 | 0.824 | 0.795 |
| *Arabis sagittata* | recovery | SSR | chr5 | 0.487 | 0.766 | 0.846 | 0.830 |

| | | | | | | | |
|---|---|---|---|---|---|---|---|
| *Arabis sagittata* | recovery | SSR | chr6 | 0.533 | 0.747 | 0.819 | 0.802 |
| *Arabis sagittata* | recovery | SSR | chr7 | 0.534 | 0.751 | 0.820 | 0.816 |
| *Arabis sagittata* | recovery | SSR | chr8 | 0.507 | 0.763 | 0.837 | 0.821 |
| *Arabis sagittata* | wilting | SSC | chr1 | 0.660 | 0.579 | 0.648 | 0.622 |
| *Arabis sagittata* | wilting | SSC | chr2 | 0.662 | 0.597 | 0.633 | 0.608 |
| *Arabis sagittata* | wilting | SSC | chr3 | 0.646 | 0.607 | 0.671 | 0.641 |
| *Arabis sagittata* | wilting | SSC | chr4 | 0.661 | 0.579 | 0.630 | 0.593 |
| *Arabis sagittata* | wilting | SSC | chr5 | 0.655 | 0.594 | 0.674 | 0.638 |
| *Arabis sagittata* | wilting | SSC | chr6 | 0.662 | 0.603 | 0.633 | 0.601 |
| *Arabis sagittata* | wilting | SSC | chr7 | 0.660 | 0.605 | 0.646 | 0.623 |
| *Arabis sagittata* | wilting | SSC | chr8 | 0.671 | 0.575 | 0.603 | 0.563 |
| *Arabis sagittata* | wilting | SSR | chr1 | 0.527 | 0.744 | 0.819 | 0.805 |
| *Arabis sagittata* | wilting | SSR | chr2 | 0.536 | 0.739 | 0.811 | 0.789 |
| *Arabis sagittata* | wilting | SSR | chr3 | 0.521 | 0.746 | 0.818 | 0.792 |
| *Arabis sagittata* | wilting | SSR | chr4 | 0.509 | 0.757 | 0.833 | 0.816 |

| | | | | | | | |
|---|---|---|---|---|---|---|---|
| *Arabis sagittata* | wilting | SSR | chr5 | 0.511 | 0.765 | 0.832 | 0.820 |
| *Arabis sagittata* | wilting | SSR | chr6 | 0.511 | 0.752 | 0.834 | 0.825 |
| *Arabis sagittata* | wilting | SSR | chr7 | 0.510 | 0.759 | 0.832 | 0.838 |
| *Arabis sagittata* | wilting | SSR | chr8 | 0.529 | 0.748 | 0.820 | 0.809 |

**Table-S7**- Details of extracted EPMs from control, wilting and recovery treatment of DeepCRE-like models including cluster assignment, positional preferences and significant similarities to JASPAR TF database. https://github.com/Abdubidopsis/Arabis_drought_transcriptome

**Table-S8:** Association of EPMs to cluster (k=25) from species and treatment wise set analyses. Nomenclature is the following for EPM detected in A. sagittata (Asag) or A. nemorensis (Anem), S0, SW and SS indicate that models were trained on a single species and on the transcriptome of control plant, wilting plants and after recovery (survivors), respectively.

| **Comparison of treatments (species-wise)** | **EPM-Cluster** |
|---|---|
| Common between AsagS0 AsagSW and AsagSS | 18, 8, 14, 25, 16, 7, 23, 22, 24, 15, 13, 1, 10, 20, 3 |
| Common between AsagS0 and AsagSW | 18, 8, 14, 25, 21, 16, 7, 23, 22, 24, 15, 13, 1, 10, 20, 3 |
| Common between AsagS0 and AsagSS | 18, 8, 14, 25, 16, 7, 23, 22, 24, 4, 15, 13, 1, 10, 20, 5, 3 |
| Common between AsagSW and AsagSS | 14, 1, 25, 6, 9, 13, 2, 15, 7, 10, 8, 16, 23, 24, 20, 19, 22, 18, 3 |
| Unique to AsagS0 | 17 |
| Unique to AsagSW | 11 |

| | |
|---|---|
| Unique to AsagSS | 12 |
| Common between AnemS0 AnemSW and AnemSS | 25, 24, 2, 7, 11, 13, 10, 22, 3, 21, 5 |
| Common between AnemS0 and AnemSW | 25, 24, 2, 7, 11, 18, 13, 10, 6, 22, 3, 21, 5, 19, 14 |
| Common between AnemS0 and AnemSS | 25, 24, 2, 8, 7, 1, 11, 13, 10, 23, 22, 3, 20, 15, 21, 5 |
| Common between AnemSW and AnemSS | 7, 25, 24, 22, 21, 10, 13, 2, 3, 11, 5 |
| Unique to AnemS0 | 9, 4 |
| Unique to AnemSW | |
| Unique to AnemSS | 12 |
| Comparison of species (treatment-wise) | EPM-Cluster |
| Common between AsagS0 and AnemS0 | 18, 8, 14, 25, 21, 7, 23, 22, 24, 4, 15, 13, 1, 10, 20, 5, 3 |
| Common between AsagSW and AnemSW | 14, 25, 6, 13, 2, 7, 10, 11, 24, 19, 22, 18, 21, 3 |
| Common between AsagSS and AnemSS | 24, 22, 13, 25, 23, 12, 8, 7, 1, 15, 2, 10, 20, 3, 5 |
| Unique to AsagS0 vs AnemS0 | 16, 17 |
| Unique to AnemS0 vs AsagS0 | 2, 11, 6, 19, 9 |
| Unique to AsagSW vs AnemSW | 1, 9, 15, 8, 16, 23, 20 |
| Unique to AnemSW vs AsagSW | 5 |
| Unique to AsagSS vs AnemSS | 4, 18, 6, 14, 16, 9, 19 |
| Unique to AnemSS vs AsagSS | 21, 11 |

**Table-S9:**Filtered annotations of EPM occurrence within the genomes of *A. nemorensis* and *A. sagitatta.* The table shows information on the chromosome number, start and end position of each EPM (column D) with further information on the strand on which the EPM is found. EPM annotation of each species are given in a separate table.
https://github.com/Abdubidopsis/Arabis_drought_transcriptome

**Table-S10:** Analyzes gene expression data linking differentially expressed genes to Gene Ontology terms with EPMs for *A. nemorensis* and *A. sagitatta*.
https://github.com/Abdubidopsis/Arabis_drought_transcriptome

**Table-S11:** Candidate transcription factor associated with genes responding differently to wilting in the two species. These transcription factors have described binding affinities that are similar to the EPM enriched in groups of differentially expressed genes.

| | | | | | EPM in *A.nemorensis* alleles | | | EPM in *A.sagittata* alleles | | |
|---|---|---|---|---|---|---|---|---|---|---|
| **Response in *A. nemorensis*** | **Response in *A. sagittata* compared to *A. nemorensis*** | **Number of genes** | **EPM_cluster** | **candidate TF** | **Total EPM number** | **Mean EPM number per gene** | **Enrichment pvalue** | **Total EPM number** | **Mean EPM number per gene** | **Enrichment pvalue** |
| Down-regulated genes | Up-regulated | 1110 | 5 | MYB-SHAQKYF1;MYS1 | 2373 | 14.92 | 4.92e-24 | - | - | - |
| Down-regulated genes | more down-regulated | 1331 | 5 | MYB-SHAQKYF1;MYS1 | 2812 | 14.49 | 2.93e-12 | - | - | - |
| Down-regulat | less down- | 1801 | 7 | HSFEA | 2709 | 3.99 | 3.98e-05 | 5439 | 6.66 | 1.13e-35 |

| | | | | | | | | | | |
|---|---|---|---|---|---|---|---|---|---|---|
| ed genes | regulated | | | | | | | | | |
| Down-regulated genes | less down-regulated | 1801 | 10 | HSFEA | 8947 | 9.28 | 1.95e-49 | 2442 | 4.25 | 6.48e-03 |
| Down-regulated genes | Up-regulated | 1110 | 11 | GATA 8, 19, 20 | 4096 | 5.88 | 6.50e-18 | 2514 | 4.50 | 4.02e-03 |
| Down-regulated genes | less down-regulated | 1801 | 13 | KAN1 | - | - | - | 6707 | 7.91 | 3.64e-26 |
| Down-regulated genes | more down-regulated | 1331 | 13 | KAN1 | 7415 | 13.96 | 4.00e-69 | - | - | - |
| Down-regulated genes | Up-regulated | 1110 | 21 | GATA 8, 19, 20 | 8944 | 11.72 | 1.51e-10 | 1533 | 4.43 | 4.64e-06 |
| Up-regulated genes | Up-regulated but to lower level | 1400 | 22 | ABI4 | 7495 | 9.45 | 6.09e-06 | 2383 | 7.52 | 1.90e-14 |
| Down-regulat | more down- | 1331 | 23 | DOF3.4/ DOF5.8 | - | - | - | 1562 | 7.27 | 3.16e-15 |

| | | | | | | | | | | |
|---|---|---|---|---|---|---|---|---|---|---|
| ed genes | regulated | | | | | | | | | |
| Up-regulated genes | Up-regulated to higher level | 2091 | 24 | GATA 8, 19, 20 | 5584 | 5.22 | 2.09e-13 | - | - | - |
| Down-regulated genes | Up-regulated | 1165 | 25 | Ethylene Response Factor family | 8202 | 15.30 | 7.79e-11 | 9946 | 17.24 | 4.42e-01 |
| Down-regulated genes | more down-regulated | 1331 | 25 | Ethylene Response Factor family | 9440 | 16.80 | 1.24e-06 | 13095 | 21.64 | 1.92e-44 |

**Table-S12:** Occurrence of EPMs and characterized TFBS in miRNA408 of *A. nemorensis* and *A. sagitatta*. https://github.com/Abdubidopsis/Arabis_drought_transcriptome

**Table-S13:** miRNA normalized expression matrix for Arabis sagittata and Arabis nemorensis in control, wilting and recovery. https://github.com/Abdubidopsis/Arabis_drought_transcriptome

### Supplementary Material

**File-S1:** Sequence alignment of miRNA408 with EPMs for *A. thaliana, A. nemorensis* and *A. sagitatta*. https://github.com/Abdubidopsis/Arabis_drought_transcriptome

**File-S2:** Sequence alignment of potential miRNA408 target with EPMs for *A. thaliana, A. nemorensis* and *A. sagitatta*. https://github.com/Abdubidopsis/Arabis_drought_transcriptome
